# Genome-wide association studies in a diverse strawberry collection unveil loci controlling agronomic and fruit quality traits

**DOI:** 10.1101/2024.03.11.584394

**Authors:** Pilar Muñoz, F. Javier Roldán-Guerra, Sujeet Verma, Mario Ruiz-Velázquez, Rocío Torreblanca, Nicolás Oiza, Cristina Castillejo, José F. Sánchez-Sevilla, Iraida Amaya

**Affiliations:** Centro IFAPA de Málaga, Instituto Andaluz de Investigación y Formación Agraria y Pesquera, (IFAPA), Málaga, Spain; Department of Horticultural Sciences, IFAS Gulf Coast Research and Education Center, University of Florida, Wimauma, Florida 33598, USA; Unidad Asociada de I+D+i IFAPA-CSIC Biotecnología y Mejora en Fresa, Málaga, Spain

**Keywords:** Fruit firmness, Genetic diversity, GWAS, KASP, Marker-assisted selection, Population structure, Strawberry

## Abstract

Strawberries are cherished for their organoleptic properties and nutritional value. However, breeding new cultivars involves the simultaneous selection of many agronomical and fruit quality traits. The strawberry germplasm collection here studied exhibited extensive phenotypic variation in 26 agronomic and fruit quality traits across three consecutive seasons. Phenotypic correlations and Principal Component Analysis (PCA) revealed relationships among traits and accessions, emphasizing the impact of plant breeding. Genetic diversity analysis on 124 accessions using 44,408 markers denoted a population structure divided into six subpopulations that still retain considerable diversity. Genome-wide association studies (GWAS) for the 26 traits unveiled 121 significant marker-trait associations distributed across 95 quantitative trait *loci* (QTL). Multiple associations were detected for fruit firmness, a key breeding target, including a prominent *locus* on chromosome 6A. The candidate gene *FaPG1,* controlling fruit softening, was identified within this QTL region. Differential expression of *FaPG1* confirmed its role as the main contributor to natural variation in fruit firmness. A Kompetitive Allele-Specific PCR (KASP) assay based on SNP AX-184242253, associated with the 6A QTL, predicts a substantial increase in fruit firmness, validating its utility for Marker-Assisted Selection (MAS). In essence, this comprehensive study provides insights into the phenotypic and genetic landscape of the strawberry collection and lays a robust foundation for propelling the development of superior strawberry cultivars through precision breeding.

**Highlight:** Several QTL controlling agronomic and fruit quality traits detected by genome wide association studies. Natural variation on *FaPG1* expression is associated with a mayor and stable QTL for fruit firmness.

## Introduction

Strawberry fruit is highly consumed and appreciated for its vibrant red color, flavor, aroma and nutritional value. These sensory attributes, and potential health benefits to consumers have contributed to the economic importance of this fruit crop, which has reached a world production of 9.5 million tons in 2022 (FAOSTAT database, 2024, http://www.fao.org). A substantial trait diversity is observed in strawberry varieties due to their elevated heterozygosity and a relatively brief breeding history spanning less than three centuries. (Darrow, 1966; Hardigan et al., 2021a). Breeding efforts in strawberry have been primarily directed towards preserving high yields, enhancing pest resistance, as well as improving fruit weight, firmness and flavor (Whitaker et al., 2017; Verma et al., 2017; Senger et al., 2022). Among fruit quality traits, increasing fruit firmness has been of maximum interest, as it influences the organoleptic quality, but most importantly the postharvest life and market value of the fruit. During strawberry ripening, fruit firmness decreases dramatically due to the induction of enzymes involved in the degradation or modification of the cell wall, including polygalacturonases (PGs), pectin methylesterases, pectate lyases, endoglucanases and expansins (reviewed in Moya-Leon et al. 2019). Extending postharvest life is essential for long-distance transportation. For instance, according to the FAOSTAT database (2024, http://www.fao.org), Spain produced 325,880 tons of strawberries in 2022, with more than 85% of this production transported to Germany and other European countries via road transportation.

In response to consumer preferences, there is a growing emphasis on enhancing fruit organoleptic and nutritional quality in current breeding programs. Therefore, many strawberry breeders worldwide address the improvement of sugar/acid balance, aroma, fruit color, and antioxidant capacity or ascorbic acid (AsA) content (Mezzetti et al., 2016; Verma et al., 2017; Senger et al., 2022). On the other hand, the main breeding objectives for growers are focused on agronomic traits related to yield (number of fruits and size), resistance to pests and diseases (Phytophthora crown rot, Fusarium wilt, Anthracnose, or powdery mildew, among many others), and key physiological traits related to plant vegetative propagation (runnering time and number of runners), flowering habit and fruit precocity.

The cultivated strawberry (*Fragaria* × *ananassa* Duchesne ex Rozier) is a complex allo-octoploid species (2n = 8× = 56) originating from the hybridization of two octoploid wild species: The South American *F. chiloensis* and the North American *F. virginiana* (Darrow, 1966; Hancock, 1999). Their octoploid genome is formed by four subgenomes derived from four diploid progenitor species. Two of these subgenomes can be traced back to the ancestors *F. vesca* and *F. iinumae*, whereas the donors of the remaining two subgenomes might be extinct and have not been clearly identified (Tennessen et al., 2014; Edger et al., 2018, 2019; Liston et al., 2020; Hardigan et al., 2021a). Genome-wide association studies (GWAS) were originally implemented in cultivated strawberry using the reference genome of the diploid *F. vesca.* Those first analyses led to successful identification of genomic regions linked to several agronomic and fruit quality traits, such as disease resistance (Pincot et al., 2018), flowering time (Hardigan et al., 2018) or volatile organic compounds (VOCs) influencing fruit aroma (Barbey et al., 2021). Since the release of the first octoploid strawberry genome sequenced with chromosome-level resolution (Edger et al., 2019), association studies in this crop have benefited from subgenome-specific detection. These studies have explored the genetic control of fruit quality traits such as color (Castillejo et al., 2020), firmness (Hardigan et al., 2021a) or VOCs (Fan et al., 2022), among others. It’s worth noting that GWAS results could be influenced by various factors, including association models, population size and structure, phenotype, and environmental factors (Gai et al., 2023).

This research aims to uncover new loci that contribute to the variation of important breeding targets and subsequently develop biotechnological tools for strawberry molecular breeding. To achieve this, we conducted GWAS using a population comprising, depending on the season, 95 to 124 different strawberry accessions from the *Fragaria* germplasm collection at IFAPA (ESP138). This population included varieties of *F.* × *ananassa* and some hybrids with *F. chiloensis* covering a broad geographic and temporal diversity, as well as a wide representation of accessions adapted to the Californian-Mediterranean climate. Accessions were genotyped using the 50K Fana Axiom platform (Hardigan et al., 2020), which was mapped on the reference octoploid genome (Edger et al., 2019). As an initial step preceding the GWAS, we analyzed the similarity among accessions, the population’s structure and their genetic diversity. Over the course of three consecutive seasons, we phenotyped this collection for 26 agronomical and fruit quality traits in order to evaluate the influence of environmental factors on these traits and identify loci that exhibit stability across years. We report previously detected and novel QTLs for important breeding targets, and also for characters used as descriptors in the UPOV examination of Distinctness, Uniformity and Stability. Among all those traits we focused our attention on a mayor QTL for fruit firmness on chromosome 6A. We searched the haploblock in linkage disequilibrium for candidate genes and, based on functional annotation and expression analyses identified *FaPG1* as the underlying gene. Furthermore, we here developed and validated a Kompetitive Allele Specific PCR (KASP) assay to assist the selection of strawberries with higher fruit firmness.

### Experimental procedures Plant material

The study has been carried out during three consecutive seasons (2018-2019, 2019-2020 and 2020-2021). The experimental population consisted of 134 strawberry accessions from the *Fragaria* germplasm collection at IFAPA, including cultivated strawberry varieties (*F.* × *ananassa*) as well as hybrids with *F. chiloensis.* The collection was selected to cover a wide genotypic, geographical and temporal diversity, with a significant representation of accessions adapted to the Californian-Mediterranean climate (Supplementary Table S1). The cultivar ’Mieze Schindler’, for which two different accessions from different sources (ACC_248 and ACC_307) were available in the collection, was used as internal control. Young plants of each genotype were obtained each season after vegetative multiplication of accessions from the IFAPA *Fragaria* germplasm collection. Six rooted plants per genotype were transplanted during November to a shaded greenhouse in Málaga, following a randomized block design. To simulate field conditions, the experimental population was cultivated in a total of 104 1x1 m containers filled with a mixture of universal substrate with river sand in a 3:1 v/v ratio. Eight plants were planted in each container at a distance of 25 cm, which is similar to the strawberry planting density in commercial fields at Huelva, Spain. Plants were irrigated automatically using a drip system three days per week, increasing to seven days during the summer period. During all seasons, plants were treated monthly against white spider mite and red spider mite, cleaned weekly of old leaves and weeds and fertilized at variable concentration according to the time of year. During the second and third seasons, plants were regularly treated against powdery mildew.

### DNA extraction, genotyping and pedigree confirmation

Genomic DNA was extracted using 40-50 mg of freeze-dried young leaf tissue following the instructions of the Omega DNA extraction kit (Bio-tek; https://www.omegabiotek.com) with minor modifications specified in Hardigan et al., 2018 and quantified in a Qubit fluorometer (Thermo Fisher Scientific). DNA from individuals were submitted to Segalab for genotyping using the Fana 50K Axiom™ array (Hardigan et al., 2020). The generated data files were analyzed with the Axiom™ Analysis Suite v.5.1.1.1 software (Thermo Fisher Scientific) following the Best Practices Genotyping Workflow described in the software documentation.

Markers were filtered for a call rate (CR) ≥ 97%, and the parameters established for polyploid species by default. SNP and indel markers assigned into the ‘Poly HighResolution’ (PHR; polymorphic with three defined clusters) or ‘NoMinor Homozygote’ (NMH; polymorphic with two defined clusters) classifications were selected for further analyses. Samples with a CR greater than 97% were obtained, although two accessions with CR of 93 and 94% were manually selected and retained.

Similarity between accessions was analyzed using the Identity By State (IBS) distance matrix using PLINK v.1.9 software (Chang et al., 2015). Accessions with higher similarity than that obtained between the two accessions of ’Mieze Schindler’ (ACC_248 and ACC_307; IBS > 99.78%) were considered propagation errors or synonyms. In order to determine one or the other situation, all genotypes were compared with data obtained for those same accessions in a previous genotyping of the IFAPA collection performed with an independent DNA extraction and the Affymetrix IStraw90K Axiom array (Bassil et al., 2015). Those accessions that turned out to be propagation errors, duplicates and/or synonyms were removed from further analyses, remaining a total of 95, 114 and 124 accessions in the first, second and third seasons.

### Genetic Diversity and Population structure

Genetic diversity analyses were performed with the 44,408 high-quality SNPs and indels selected from the Affymetrix 50K Fana Axiom array (Hardigan et al., 2020) in 124 accessions of the experimental population using different methods. Principal Component Analysis (PCA) was implemented using TASSEL v.5.0 (Bradbury et al., 2007). The population structure of the experimental population was analyzed using 44,002 SNPs (w/o indels) and the software STRUCTURE v.2.3.4 (Pritchard et al., 2000). Two to 20 sub-populations were evaluated with an admixture model, 10,000 burn-in steps, 100,000 Markov-Chain Monte Carlo (MCMC) steps and 10 replicates per K-values. This analysis was carried out at the facilities of the Supercomputing and Bioinnovation center of the University of Malaga (https://scbi.uma.es) using StrAuto (Chhatre & Emerson, 2017). The optimal number of subpopulations (K value) was established using STRUCTURE HARVESTER (Earl & vonHoldt, 2012) and the Evanno method (Evanno et al., 2005). Sample orders were calculated using CLUMPP v. 1.1.2 (Jakobsson & Rosenberg, 2007) and cluster results were visualized using Distruct v.1.1 (Rosenberg, 2004). Neighbor-joining phylogenetic tree was constructed with TASSEL v.5.0 (Bradbury et al., 2007) from the 1-IBS distance matrix. The dendrogram obtained was drawn and visualized with Fig Tree v.1.4.4 (http://tree.bio.ed.ac.uk/software/figtree/). Finally, the pairwise fixation index (FST) was calculated using the Hierfstat v0.5-7 tool for *R* (Goudet 2005) with data obtained from the STRUCTURE, to know the degree of population structuring.

### Phenotypic evaluation

Phenotypic characterization for a total of 26 morphological, physiological and fruit quality traits was carried out during three consecutive years. Trait data collection was carried out in the greenhouse weekly, during the optimum of plant and/or fruit development. Traits were noted by the letters FL if they relate to flowers, LE if they relate to leaves, PL for plant-related, RU for runners, and FR for fruit-related, followed by the name of the traits or by an abbreviation of it. Reference scales established for variety protection by UPOV (https://cpvo.europa.eu/sites/default/files/documents/fragaria_3.pdf) were used for 12 traits.

Powdery mildew natural infection on the plant (no inoculation performed) was evaluated on a regular basis from November until August only during the first season, using a 1-5 rating scale based on the presence of powdery mildew symptoms on the leaf: (0) healthy plant with 0% presence, (1) <10%, (2) 11-25%, (3) 26-50% or (4) >50% leaf area affected.

Besides using the UPOV scale, external fruit color was measured in a total of eight fruits per accession with a chromameter (Chroma Meter CR 410, Konica-Minolta) using CIELAB color space values *a\** (green–red spectrum), *b\** (blue–yellow spectrum), and *L\** (brightness– darkness).

Fruit firmness was evaluated using a penetrometer (Facchini Fruit Pressure tester FT 02) with a 3.5 mm probe in eight fruits per genotype taking two measurements on each fruit.

For quality traits evaluated in processed fruit, at least 25 commercially mature fruits of each genotype were harvested weekly, at approximately the same time of the morning, and during the production peak of the season. These fruits were immediately cut, frozen in liquid nitrogen and stored at -80 °C. Fruits of each accession were divided into three biological replicates consisting of a minimum of eight fruits. The set of fruits from each replicate was powdered in liquid nitrogen using a coffee grinder and stored at -80 °C until analysis. The first fruits produced by the plant were discarded because they were not representative of each variety, as well as misshaped fruits. Fruit acidity (citric acid equivalents: g citric acid/100g FW) was measured with an automatic titrator (TitroLine easy, Schott Instruments®, GmbH) by diluting 1 g of fruit powder from each replicate in 100 mL of distilled water. Titrations were carried out to a final pH of 8.1 using 0.01 N NaOH. Soluble solids content (SSC) in Brix degrees was quantified using a digital refractometer (ATAGO™ Brix digital refractometer PR-32α) and a drop of blended fruit from the three replicates. Ascorbic acid content (mg AsA/100 g FW) was quantified spectrophotometrically using the protocol of Fenech et al., (2021) with modifications: approximately 100 mg of the fruit powder from each replicate was homogenized in 1 mL of cold extraction buffer (3% metaphosphoric acid, 1 mM EDTA) and centrifuged at 14,000 rpm for 20 min at 4 °C. The supernatant was filtered through a 0.45 µm nylon membrane. Absorbance measurements were carried out at 265 nm in the plate reader on UV-transparent 96-well plates (Greiner UV-Star 96-Well Microplate) after 30 min incubation of 20 µL of the filtered sample with 100 µL of phosphate buffer (0.2 MKH2PO4). After the measurement, 5 µL of the ascorbate oxidase enzyme (40 U/mL in phosphate buffer) was added to each well, mixed and incubated at room temperature for 20 min, and the absorbance was measured again. The AsA content was calculated by interpolating the difference of the absorbance on a standard curve with known concentrations of AsA (Sigma).

Correlations for each trait between the three years, and correlations between the log-transformed data (log10) of the 25 traits measured in the third season (2020-2021) were calculated using Pearson correlation in *R* version 4.3.2 using ‘stats’ and ‘corrplot’ packages. Principal component analysis of 124 accessions of the third season was performed with the log-transformed data (log10) of the 25 traits in *R* using ‘FactoMineR’ and ‘Factoextra’ packages. Boxplots of fruit firmness across subpopulations were calculated using the *R* package ‘ggpubr’.

### Genome-Wide Association Study (GWAS)

GWAS was performed with the genotypic data from the 50K Fana Axiom array (Hardigan et al., 2020) and phenotypic data from each season separately. Missing genotype data was imputed with Beagle v.4.0 (Browning and Browning, 2007) obtaining a total of 124 accessions genotyped with 44,196 markers mapped to the reference ’Camarosa’ v1.0. a2 genome (Liu et al., 2021). For phenotypic data, the mean of the biological replicates for each accession in each year was used. Those traits that deviated from normality were transformed using the COX- BOX transformation (Box & Cox, 1964) only when transformation improved the normality. GWAS was conducted in the *R* package GAPIT v.3 (Lipka et al., 2012; Wang & Zhang, 2021) using the GLM, MLM, FarmCPU and BLINK analysis models (Price et al., 2006; Yu et al., 2006; Liu et al., 2016; Huang et al., 2019). All associations with p-value < 0.05 after the False Discovery Rate (FDR) control procedure were considered significant. Those significant markers that were within a distance of less than 5 Mb were considered a single QTL. To take into account the population stratification, the genomic relationship matrix (G) was generated using the VanRaden algorithm (VanRaden, 2008) and included as random effect, while the top three principal components, explaining a 26.1% of the variance, were used as covariates in the GWAS model. Haplotype blocks were computed with ‘SnpStats’ package in R using ld function to calculate pairwise LD (D’) and plotted with ld.plot function from the ‘gaston’ package.

### *In silico* candidate gene identification

Genes associated with the positions of the significant SNPs were obtained by searching for the position of the significant SNP marker in the *Genome Database for Rosaceae* (GDR; https://www.rosaceae.org/). The gene located at that specific position was identified, or in cases where no gene was found, the flanking genes to it. Subsequently, a BLAST analysis (Basic Local Alignment Search Tool) was performed to determine the function assigned to each of the associated genes using the ’Camarosa’ reference genome v1.0 (Edger et al., 2019) with the new a.2 annotation (Liu et al., 2021).

### RNA isolation and gene expression analysis

We generated two pools of fruits selecting 30 accessions contrasting in fruit firmness among the 124 accessions used in the third-season (2020-2021) GWAS. Each pool in triplicate consists of an equivalent amount of fruit tissue stored at -80°C from each of the three biological replicates of 15 accessions. We isolated total RNA from the three biological replicates from each pool of 15 accessions (high fruit firmness vs. low fruit firmness) using the Plant/Fungi Total RNA Purification Kit (Norgen Biotek). Prior reverse transcription, RNA was treated with DNAse Turbo (Invitrogen) following to the manufacturer’s instructions. A total of 800 ng of RNA was used for retrotranscription using the High-Capacity cDNA Reverse Transcription Kit (Applied biosystems by Thermo Fisher Scientific). Primers for quantification of *FaPG1* by qRT-PCR, FaPG1_F: GCTCCTGGTGACTTTGATGT and FaPG1_R: ACTCTACTTGGCGTTGTTGC, were described in López-Casado et al., 2023. Subgenome- specificity of primers for *FaPG1* was confirmed by comparison to the other PG homoeologs and the most similar homologs retrieved from the GDR database (https://www.rosaceae.org/). qRT-PCRs were conducted in a CFX96™ Real-Time System (Bio-Rad) using 2 µL of a 1:5 dilution of the cDNA, 250 nM of each specific primer and the TB Green Premix Ex Taq (Takara) in a final volume of 10 µL. Three technical replicates were used for each biological replicate. The PCR program involved an initial denaturation at 95°C for 30 s, followed by 39 cycles of denaturalization at 95°C for 10 s, and annealing at 60°C for 25 s. Relative expression of the *FaPG1* gene was calculated by the 2^-ΔΔCT^ method, using *FaGAPDH* and *FaDBP* as reference genes (Muñoz-Avila et al., 2022) and the High-Firmness pool as the reference sample. Statistical analysis was performed using GraphPad Prism 8.0 software (San Diego, California, USA). The normality of data distribution was evaluated by Kolmogorov–Smirnov and then analyzed by t test.

### Kompetitive allele specific PCR (KASP) assay

A KASP assay for AX-184242253, one of the significant SNPs detected on chromosome 6A, was designed using PolyOligo (https://github.com/MirkoLedda/polyoligo) and the ‘Camarosa’ v.1.0 reference genome. To further reduce the possibility of off-targets, primers were manually adjusted for increased sub-genome specificity (FIRM6A-KASP_REF: GAAGGTCGGAGTCAACGGATTattgccacaaatgcaagagG, FIRM6A-KASP_ALT: GAAGGTGACCAAGTTCATGCTattgccacaaatgcaagagA, and FIRM6A-KASP_COM: GCTAAAACATGACAACATCATCTTAC). KASP assays were performed on a CFX96 Real-Time thermal cycler (Bio-Rad) using the KASP-TF Master Mix (LGC Genomics, Teddington, UK). The PCR program consisted of an activation at 94°C for 15 min, 10 touchdown cycles of 94°C for 20 s and 61-56°C for 1 min (decreasing 0.5°C per cycle), followed by 28 cycles of 94°C for 20 s and 56°C for 1 min, and plate reading after one step of 1 min at 37°C. One or two recycling programs consisting of three additional cycles were necessary for cluster improvement and/or amplification of some accessions. Statistical analyses were performed using Student’s t-test implemented in GraphPad Prism version 8.0 software (San Diego, California USA).

## Results

### Genetic diversity and population structure

After the analysis of similarity using the IBS calculations and pedigree confirmation of the 134 accessions of the experimental population, 10 of them were removed from the study due to possible propagation errors (five accessions), duplicates and/or synonyms (Supplementary Table S2). Among synonyms, accessions ‘Revada’, ‘Georg Soltwedel’, ‘Africa’, and ‘Gorella White Pulp’ were considered synonyms and only ‘Africa’ (ACC_256) was retained in the studies. Similarly, ‘Victorian Nameless’ and ‘Surprise des Halles’ were considered synonyms and only ‘Surprise des Halles’ (ACC_266) was retained. Therefore, genetic diversity, population structure and FST index were analyzed using a total of 124 accessions and 44,408 high quality SNPs and indels.

Evaluation of population structure using PCA resulted in the two first principal components (PC) explaining 16 and 5% of the variance (Fig. 1A). PCA organized the experimental population by time and geographic adaptation, with the most modern Californian- Mediterranean varieties at the left side (in red color) and another distinct group including the oldest European varieties adapted to northern territories (in blue color) at the right side. The rest of varieties were distributed between these two groups according to time and climatic/geographical adaptation. Recently developed hybrids between Californian varieties and *F. chiloensis* (highlighted in green) clustered in the lower and central part of the graph.

**Fig. 1.**
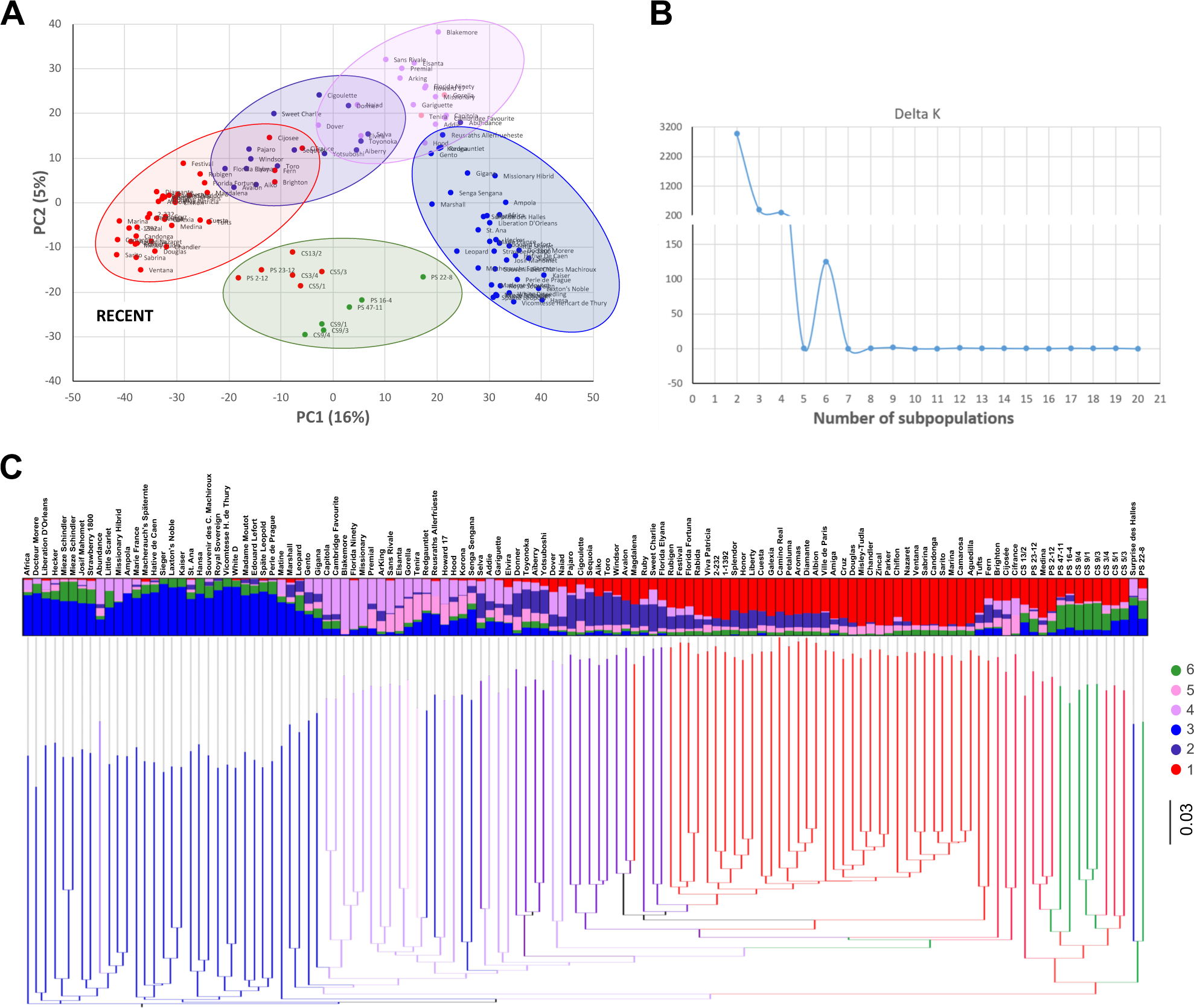
Genetic diversity and population structure of 124 strawberry accessions. (A) Principal Component Analysis (PCA) based on 44,002 SNPs describing the relationships among accessions using TASSEL (Bradbury et al., 2007). Hybrids with *F. Chiloensis* (clustering on the right side of panel C) have been enclosed on a green circle. (B) Determination of the number of subpopulations using the Delta K method (Evanno et al., 2005). (C) Dendrogram of similarity by Neighbour-Joining. Accessions have been color-coded according to their membership to the 6 STRUCTURE subpopulations. NJ-tree implemented in TASSEL (Bradbury et al., 2007). For each accession, the color bar represents the proportion of ancestry using K=6 ancestral groups inferred with the STRUCTURE program (Pritchard et al., 2000).

To assess the level of genetic stratification we used the software STRUCTURE, which showed a diversification of the strawberry collection into six subpopulations (Fig. 1B and 1C). The largest subpopulation includes classic Californian cultivars ‘Chandler’ and ‘Camarosa’ and more recent Californian/Mediterranean/Floridian varieties, as ‘Candonga’ or ‘Fortuna’ (subpopulation 1; Fig. 1C). The second subpopulation (in dark purple in Fig. 1C), contains a considerable level of admixture and is formed mainly by older Californian/Floridian varieties, such as ‘Pajaro’, ‘Toro’ or ‘Aiko’, and some Japanese accessions such as ‘Aiberry’ or ‘Toyonoka’. The oldest European accessions from northern territories belong to a third subpopulation (in blue in Fig. 1C), which includes ‘Laxtońs Noble’ and ‘Vicomtesse’. Two subpopulations act as a transition between the blue and purple (subpopulations 4 and 5; Fig. 1C). In subpopulation 4 (light purple) stand out some varieties defined by Darrow (1966) as "large-fruited strawberries best adapted to tropical climates" (‘Blakemore’, ‘Florida ninety’, ‘Missionary’) and classic North-American varieties as ‘Howard 17’. Subpopulation 5 (in pink) contains only two accessions from the Netherlands, ‘Tenira’ and ‘Gorella’. Finally, the population structure analysis discriminated a sixth subpopulation consisting of varieties introgressed with *F. chiloensis* (in green; Fig. 1C) that cluster with other related hybrids belonging to subpopulation 1.

Despite the identification of six subpopulations, the degree of stratification was scarce, and subpopulations overlapped with each other in the dendrogram. This was corroborated by a fixation index value (FST) of 0.1343 for the experimental population, indicating a limited differentiation between subpopulations and the absence of clearly defined clusters in the collection.

### Phenotypic variation of agronomic and fruit quality traits within the strawberry germplasm collection

The phenotypic evaluation of the experimental population was carried out during three consecutive seasons for morphological, physiological and fruit quality traits. In the three seasons, a large phenotypic variation was observed among accessions for all evaluated traits (Table 1). This variation was consistent among the three years for most of the traits, with a medium-high correlation coefficient (0.40 - 0.80) between seasons, except for the green color of petals, which was not significant between any season, suggesting that it is a trait with a strong environmental influence (Table 1). For flower diameter and fruit number, high correlations were only observed between the first and second or the second and third years, respectively. High correlations among the three seasons were observed for runnering time, number and color, indicating a low environmental influence on these traits. Fruit firmness ranged from 100 to 340 g during the three seasons and showed a 0.87 correlation between the second and third seasons, but a low correlation with the first season. In contrast, the width of fruit band without achenes displayed high correlations between the three years, again indicating a lower environmental influence.

**Table 1.**
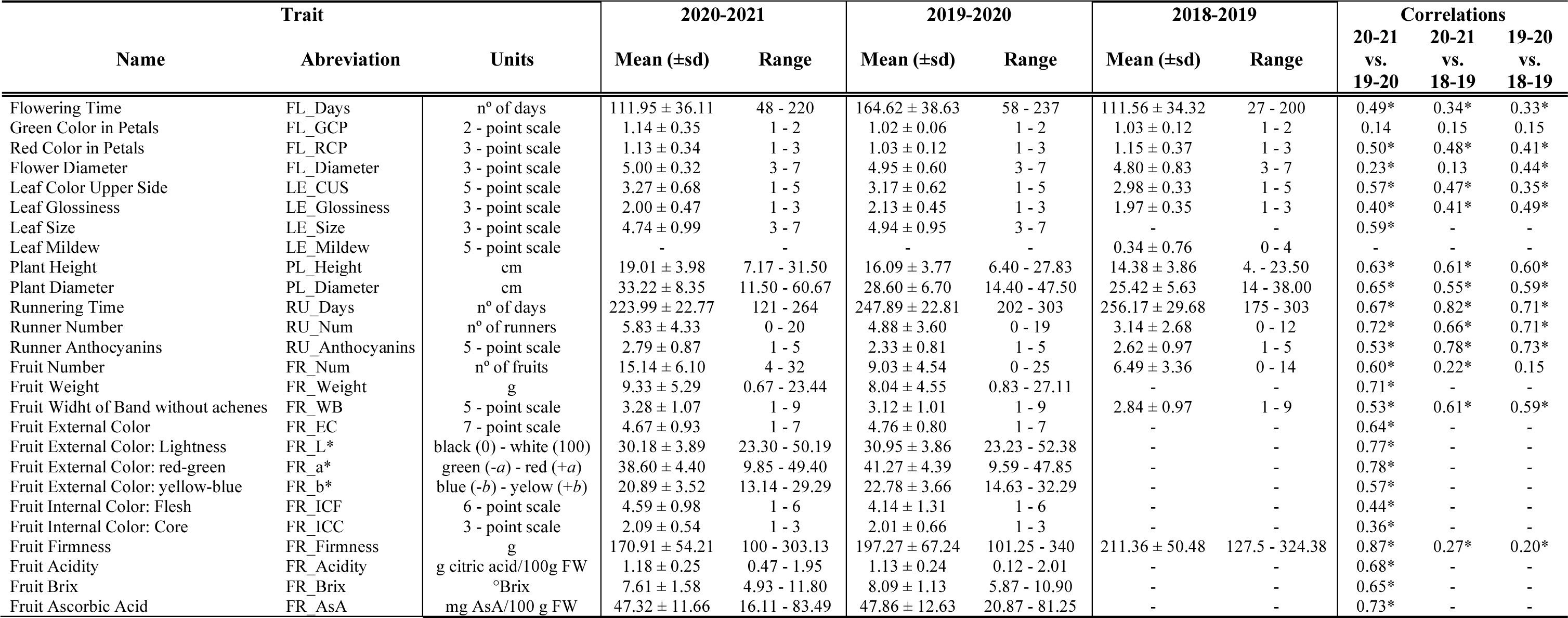
Summary of the traits evaluated during the three seasons. Means (± standard deviation), phenotypic range in the population and correlations between the three seasons for each character. *Significant correlations (p-value < 0.05) by Pearson correlation.

To identify relationships between the traits, we analyzed Pearson correlations between them using data from the third season (2020-2021; Fig. 2A; Supplementary Table S3). A total of 43 positive and 35 negative significant correlations were observed, of which 11 were considered high (± 0.60 - ± 0.80), 14 were moderate (± 0.40 - ± 0.60) and most of the significant correlations were therefore low (< ± 0.40). In general, we observed medium-high positive correlations between traits related to plant vigor (LE_Size, PL_Height, PL_Diameter) and fruit or runner production (FR_Num, FR_Weight and RU_Num). Among them, the highest significant correlations were observed between leaf size with plant height and plant diameter (LE_Size / PL_Height: +0.67 and LE_Size / PL_Diamenter: +0.66), and between plant height and plant diameter (PL_Height / PL_Diameter: +0.78). On the other hand, runnering time (RU_Days) was negatively correlated with four traits: PL_Height, PL_Diameter, LE_Size and number of runners (RU_Num), with the latter correlation standing out with a value of -0.72.

**Fig. 2.**
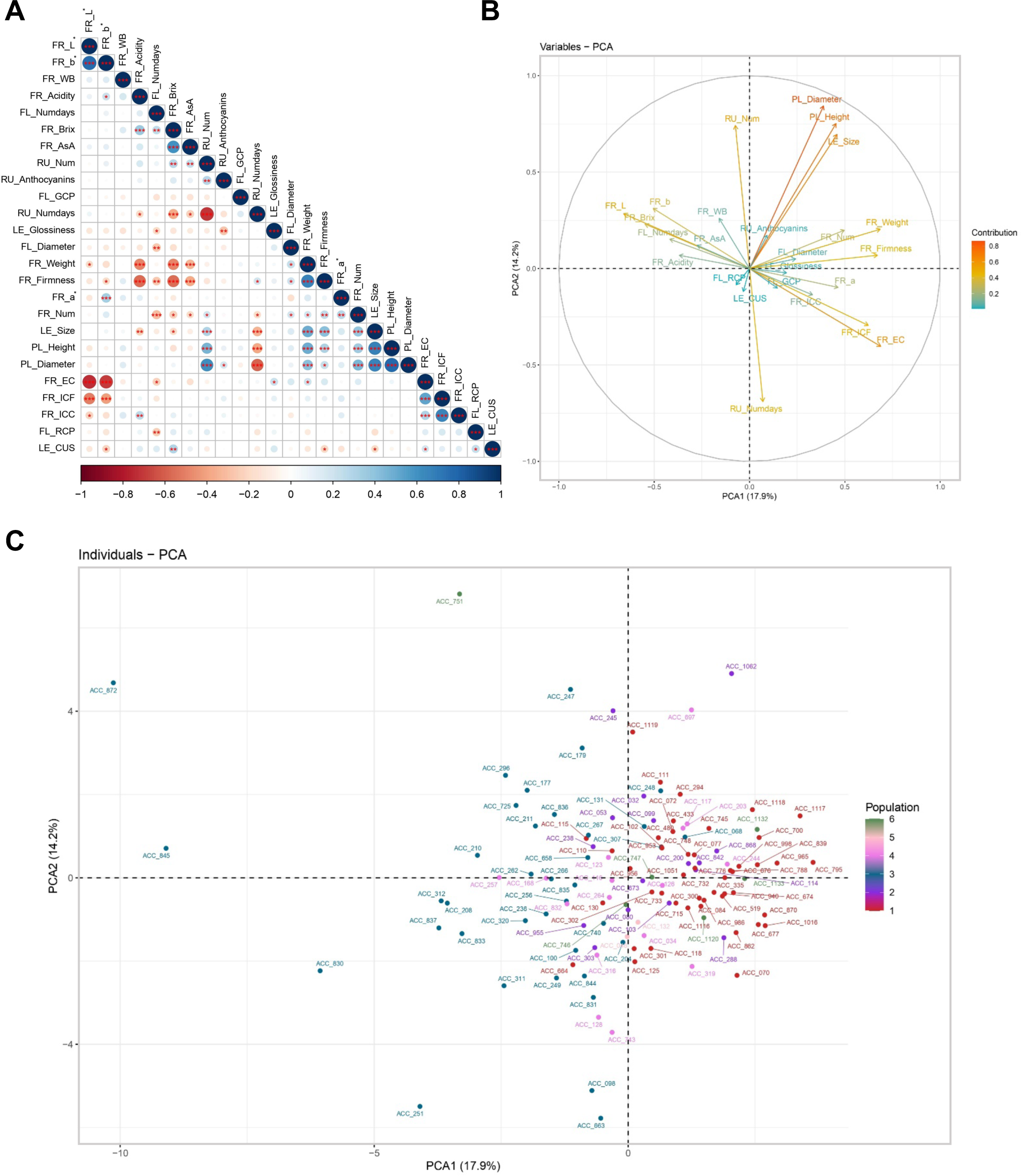
Agronomic and fruit quality traits in the strawberry collection evaluated in the 2020- 2021 season. (A) Correlation matrix among the 25 strawberry traits. Each circle indicates Pearson correlation coefficient value (r) in a scale from red (negative correlation) to blue (positive correlation). * p-value ≤ 0.05; ** p-value ≤ 0.01; *** p-value ≤ 0.001. Principal component analysis (PCA) of strawberry accessions showing trait vectors (B) and accession distribution along the first two principal component axes (C). Strawberry accessions are color- coded in C according to their population structure.

Fruit traits related to color also showed medium and high correlations. We observed positive correlations between the two internal fruit color traits (FR_ICF / FR_ICC: +0.65), and between fruit brightness and yellow hue (FR_L* / FR_b*: +0.77). Negative correlations have been observed between external color (visual scale) and brightness (FR_EC / FR_L*: -0.74) or yellow hue (FR_EC / FR_b*: -0.69). This is to be expected since higher values of L* or b* are associated with lighter fruit colors (Zorrilla et al., 2011). A strong positive correlation (0.66) was observed between FR_Firmness and FR_Weight. Strikingly, FR_Firmness was negatively correlated with fruit acidity (FR_Acidity), FR_Brix and FR_AsA content (-0.57, -0.55 and - 0.38, respectively). A low positive correlation (0.37) was observed between FR_Brix and FR_Acidity. As previously reported, FR_Weight was negatively correlated with FR_Brix, FR_Acidity and FR_AsA (-0.49, -0.48 and -0.34, respectively; Stevens et al., 2007; Wada et al., 2020; Hummer et al., 2022; Muñoz et al., 2023). Albeit weakly, fruit number was also negatively correlated with FR_Brix and FR_AsA.

A PCA of phenotype data supported the observed relationships (Fig. 2B and C). In example, traits related to plant vigor cluster together in the loadings plot and had an important contribution to the separation of strawberry accessions across PC2, together with runner number and runnering time, the latter in the opposite direction. ACC_1062, at the right top of the plot, is an example of the most vigorous genotypes. Similarly, strawberry accessions were separated across PC1 based on fruit color, fruit firmness and weight, which were also great contributors. Accessions 845, 872 and 751, on the left side, produce white or light-red fruits while accessions with the darkest fruits appeared on the opposite side of the PCA. These three traits clearly separated older from recent accessions across PC1 in accordance to their population structure. For instance, fruit firmness in the third season showed significant differences between the six subpopulations, indicating a strong selection over this trait across breeding history (Supplementary Fig. S1). Another important contributor to PC1 but in an opposite direction to fruit firmness was FR_Brix, in accordance with the significant negative correlation observed between them.

### Identification of Phenotype-Genotype associations for agronomic and fruit quality traits

Association analyses have been carried out for each year separately with the GLM and MLM single-locus test models and with the FarmCPU and BLINK multi-locus test models. A total of 121 significant marker-trait associations (FDR-adjusted p-value < 0.05) were detected in 19 of the 26 evaluated traits, distributed in 95 QTL, in the three seasons and with the four models used (Table 2; Supplementary Table S4). The number of associated SNPs varied largely between seasons and models (Table 3). In the first, second and third seasons, 9, 36 and 105 marker/traits associations were detected, respectively. Regarding GWAS models, the majority of associations were detected with FarmCPU and BLINK (51 and 56), next 36 using GLM and only 7 with MLM. In general, SNPs associated with a trait detected with one model were also detected above the background with other models albeit not being significant.

**Table 2.** Number of significant SNPs and QTL detected in the GWAS for the 19 traits in any of the three seasons.

| **Trait** | **Significant SNPs** | **Significant QTL** |
| --- | --- | --- |
| PL_Diameter | 2 | 2 |
| RU_Anthocyanins | 1 | 1 |
| RU_Days | 10 | 8 |
| RU_Num | 10 | 8 |
| LE_CUS | 4 | 4 |
| LE_Glossiness | 1 | 1 |
| LE_Mildew | 1 | 1 |
| FL_RCP | 10 | 8 |
| FL_Diameter | 1 | 1 |
| FR_Num | 12 | 12 |
| FR_Weight | 4 | 4 |
| FR_WB | 1 | 1 |
| FR_L* | 6 | 5 |
| FR_a* | 13 | 12 |
| FR_b* | 4 | 4 |
| FR_EC | 4 | 4 |
| FR_Acidity | 2 | 2 |
| FR_Brix | 10 | 10 |
| FR_Firmness | 25 | 7 |
| *Total* | *121* | *95* |

**Table 3.** Summary of significant SNPs detected in each season and with each of the four GWAS models.

| **Model** | **2020-2021** | **2019-2020** | **2018-2019** | **Total SNPs /model** |
| --- | --- | --- | --- | --- |
| **GLM** | 29 | 6 | 1 | 36 |
| **MLM** | 6 | 1 | 0 | 7 |
| **FarmCPU** | 31 | 17 | 3 | 51 |
| **BLINK** | 39 | 12 | 5 | 56 |
| *Total SNPs/season* | *105* | *36* | *9* | *150* |

No QTL was detected for flowering time (FL_Days), petal greening (FL_GCP), leaf size (LE_Size), plant height (PL_Height), fruit internal color (FR_ICF and FR_ICC) or ascorbic acid (FR_AsA). Out of the total 95 QTL, only six were detected in more than one season (Supplementary Table S4). Only one QTL was detected in one individual season for the presence of anthocyanins in the runner (RU_Anthocyanins), for leaf glossiness (LE_Glossiness), leaf powdery mildew (LE_Mildew), flower diameter (FL_Diameter) and for presence of an achene-free band on the fruit (FR_WB; Fig. 3 and Supplementary Fig. S2). Two independent QTL on chromosomes 6D and 7A were detected for fruit acidity (FR_Acidity; Fig. 4A) and another two for plant diameter (PL_Diameter; Supplementary Fig. S3). Four QTL were detected for external fruit color evaluated according to the UPOV visual scale (FR_EC; Fig. 4B), the blue-yellow value of fruit external color (FR_b*; Supplementary Fig. S4), leaf color (LE_CUS; Supplementary Fig. S5) and for fruit weight (FR_Weight; Supplementary Fig. S6). A total of five QTL were detected for the external fruit brightness (FR_L*; Supplementary Fig. S7). In contrast, seven or more QTLs were detected for the remaining seven traits (Table 2), indicating a more complex genetic architecture for those traits.

**Fig. 3.**
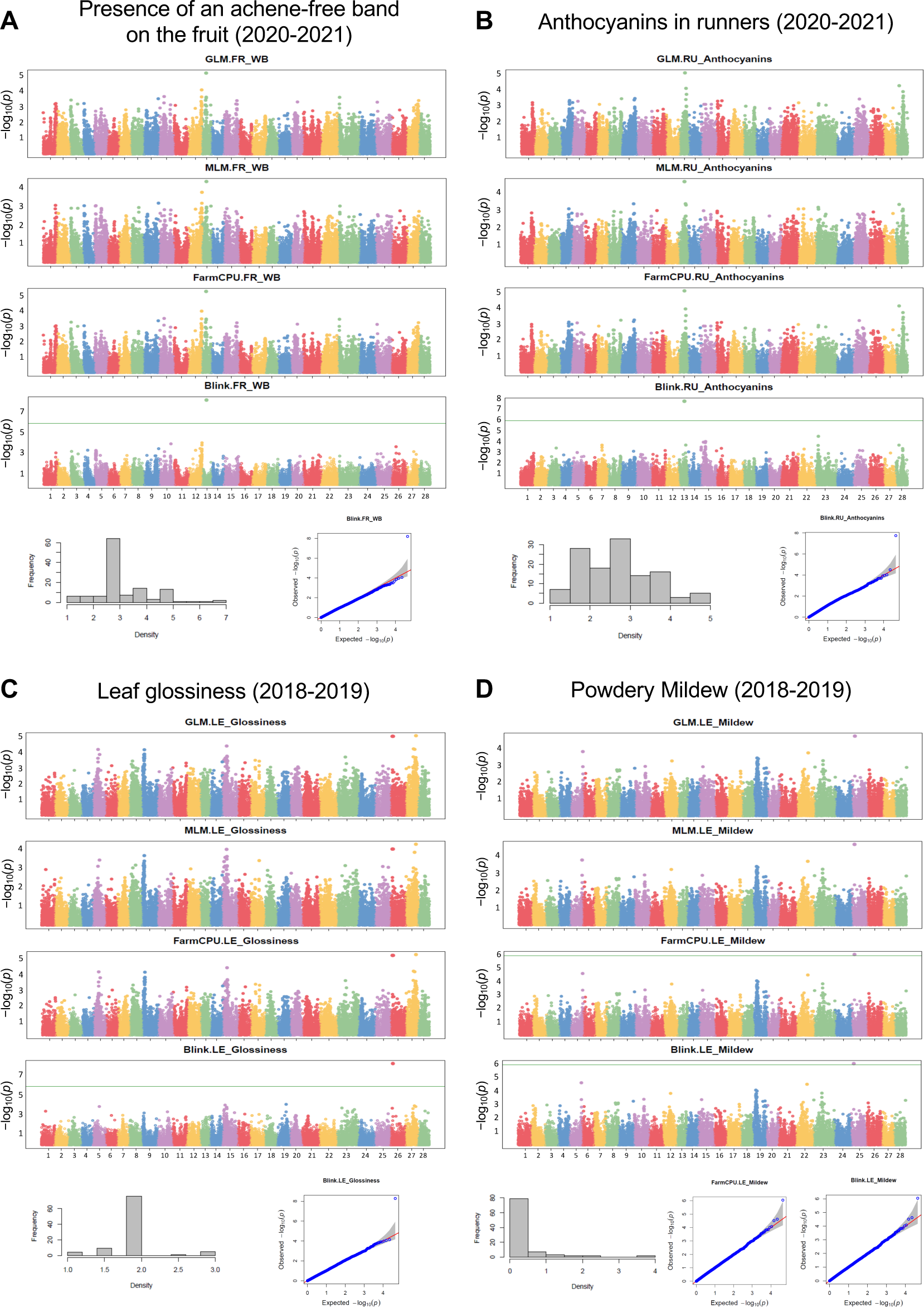
Significant GWAS associations for presence of an achene-free band on the fruit (A), anthocyanins in runners (B), leaf glossiness (C), and leaf Powdery Mildew (D). Manhattan plots of −log10 P versus chromosomal position for the four models, phenotypic distribution of traits, and QQ plots of the model(s) with the highest number of associations. Different chromosomes are shown in different colors, which follow the order of ’Camarosa’ v.1.0 (Edger et al., 2019): chromosome 1 (1-1) to chromosome 28 (7-4). Solid and dashed green horizontal lines represent the significance threshold following the Bonferroni correction method (-log10 (0.01/total SNPs) and (-log10(0.05/total SNPs), respectively).

**Fig. 4.**
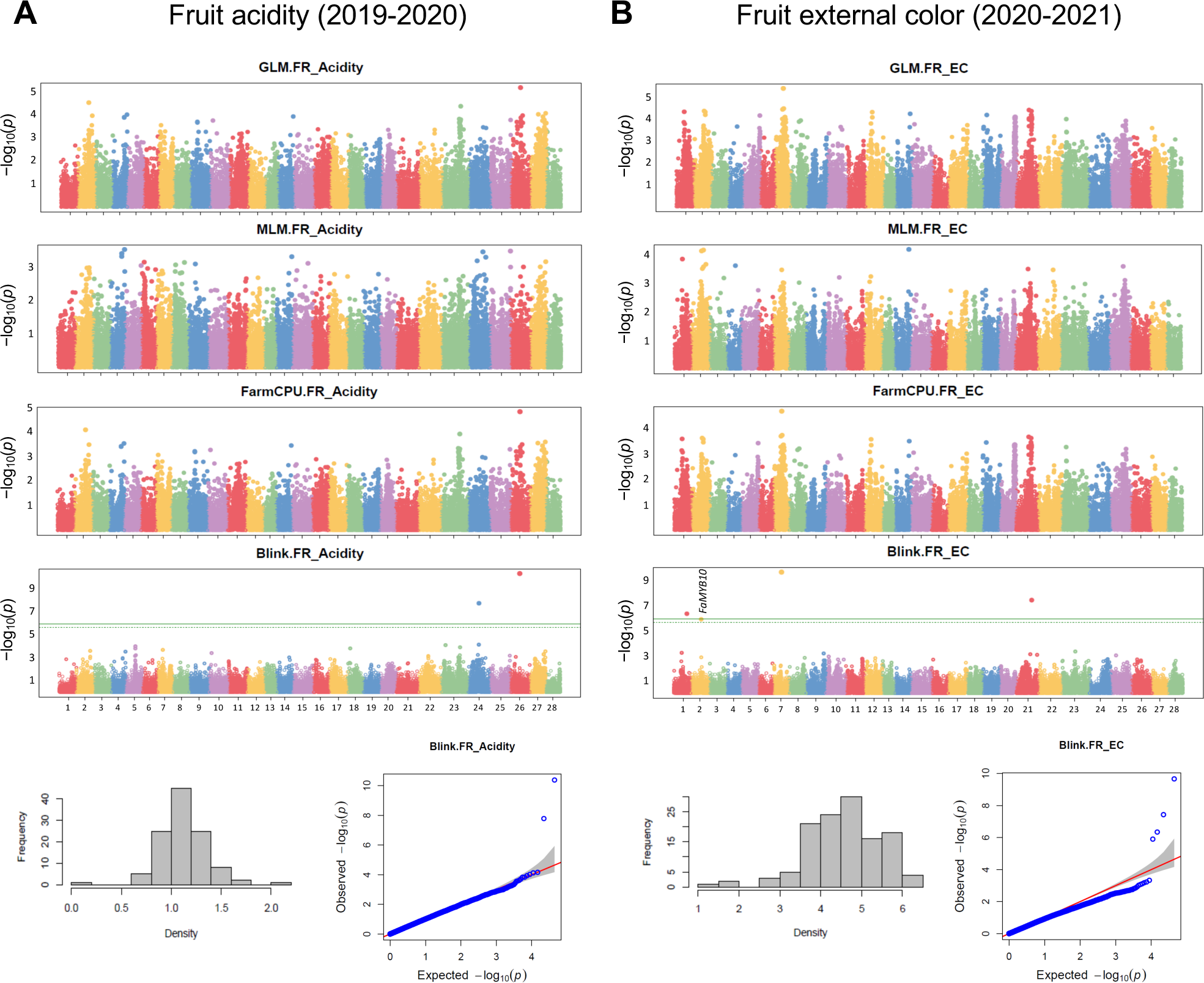
Significant GWAS associations for fruit acidity (A) and external color (B). Manhattan plots of −log10P versus chromosomal position for the four models, phenotypic distribution of traits, and QQ plots of the model with significant associations. Different chromosomes are shown in different colors, which follow the order of ’Camarosa’ v.1.0 (Edger et al., 2019): chromosome 1 (1-1) to chromosome 28 (7-4). Solid and dashed green horizontal lines represent the significance thresholds following the Bonferroni correction method (-log10 (0.01/total SNPs) and (-log10(0.05/total SNPs), respectively).

Runnering is an important trait for plant breeders and producers, and our results support a complex architecture, as eight QTL were detected for both runnering time (RU_Days; Fig. 5A) and number of runners (RU_Num; Fig. 5B). One QTL for RU_Days on chromosome 2B and two for RU_Num on 4A and 5A were detected in two seasons (Fig. 5 and Supplementary Table S4). Interestingly, the QTL on chromosome 5A has pleiotropic effects on both related traits, and i.e. the common SNP AX-184418817 delayed runnering by 18 days and decreased runner number by 1.6 - 3.3. In contrast to runnering, the presence of red coloration on petals (FL_RCP) is not currently a breeding target, and the genetic control of this trait has not been studied previously in strawberry. We have detected 10 associations in eight chromosomal regions considering all three seasons (Supplementary Fig. S8 and Table S4), being the QTL on chromosome 3B common in the first and second seasons.

**Fig. 5.**
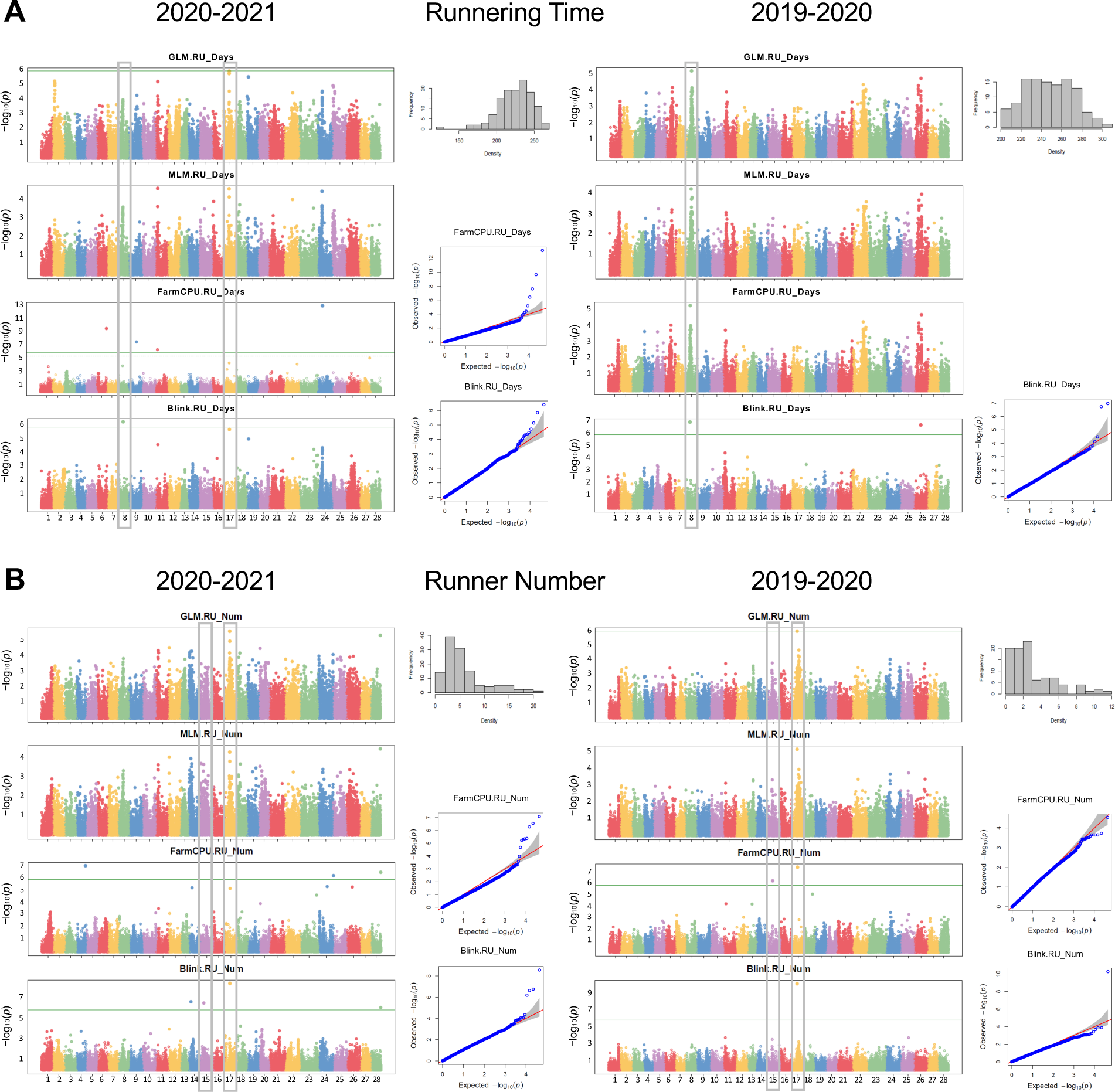
Significant GWAS associations for runnering time (A) and runner number (B) in 2020- 2021 and 2019-2020. Manhattan plots of −log10P versus chromosomal position for the four models, phenotypic distribution of traits, and QQ plots of the model(s) with the highest number of associations. Different chromosomes are shown in different colors, which follow the order of ’Camarosa’ v.1.0 (Edger et al., 2019): chromosome 1 (1-1) to chromosome 28 (7-4). Solid and dashed green horizontal lines represent the significance thresholds following the Bonferroni correction method (-log10 (0.01/total SNPs) and (-log10(0.05/total SNPs), respectively). Significant SNPs detected in two seasons or in the two related traits are highlighted.

Crop production or yield is considered as the most important breeding target and depends on both fruit number and weight. While only four QTL were detected for FR_Weight during the third season, a total of 12 QTL with small effects were detected for fruit number (FR_Num) in the 2019-2020 or the 2020-2021 seasons (Supplementary Fig. S9 and Table S4). None of them was detected in two seasons. A total of 12 QTL were also detected for the green-red value of fruit external color (FR_a*), with two of them, on chromosomes 1B and 6A, being detected in the last two seasons (Supplementary Fig. S10 and Table S4). Remarkably, the QTL on 1B, at position 15.39 Mb, was shared with other related color traits (FR_L* and FR_EC), and one of the leading SNPs, AX-184967514 (Fig. 4B; Supplementary Table S4 and Table S5), is located at the transcription factor *FaMYB10*, a positive regulator of anthocyanin biosynthesis and strawberry fruit color (Medina-Puche et al., 2014; Castillejo et al., 2020).

Our GWAS also showed a complex genetic architecture for soluble solids content (FR_Brix), with one and nine significant associations in the second and third season, respectively (Supplementary Fig. S11; Supplementary Table S4). Fruit firmness was the trait with the highest number of significant associations, 25 SNPs distributed in a total of seven QTL on chromosomes 1A, 2B, 3A, 5A, 5D, 6A and 7A. Only one QTL was detected in the second season and the rest in the third (Fig. 6; Supplementary Table S4). QTL on chromosomes 3A and 6A were each characterized for having 10 significant SNPs in an interval of 1.2 Mb and 4.3 Mb, respectively. Significant SNPs on 6A were detected with the four GWAS models. It is also worth noting that the effects on fruit firmness were up to 30 g for each of these two QTL (Supplementary Table S4).

**Fig. 6.**
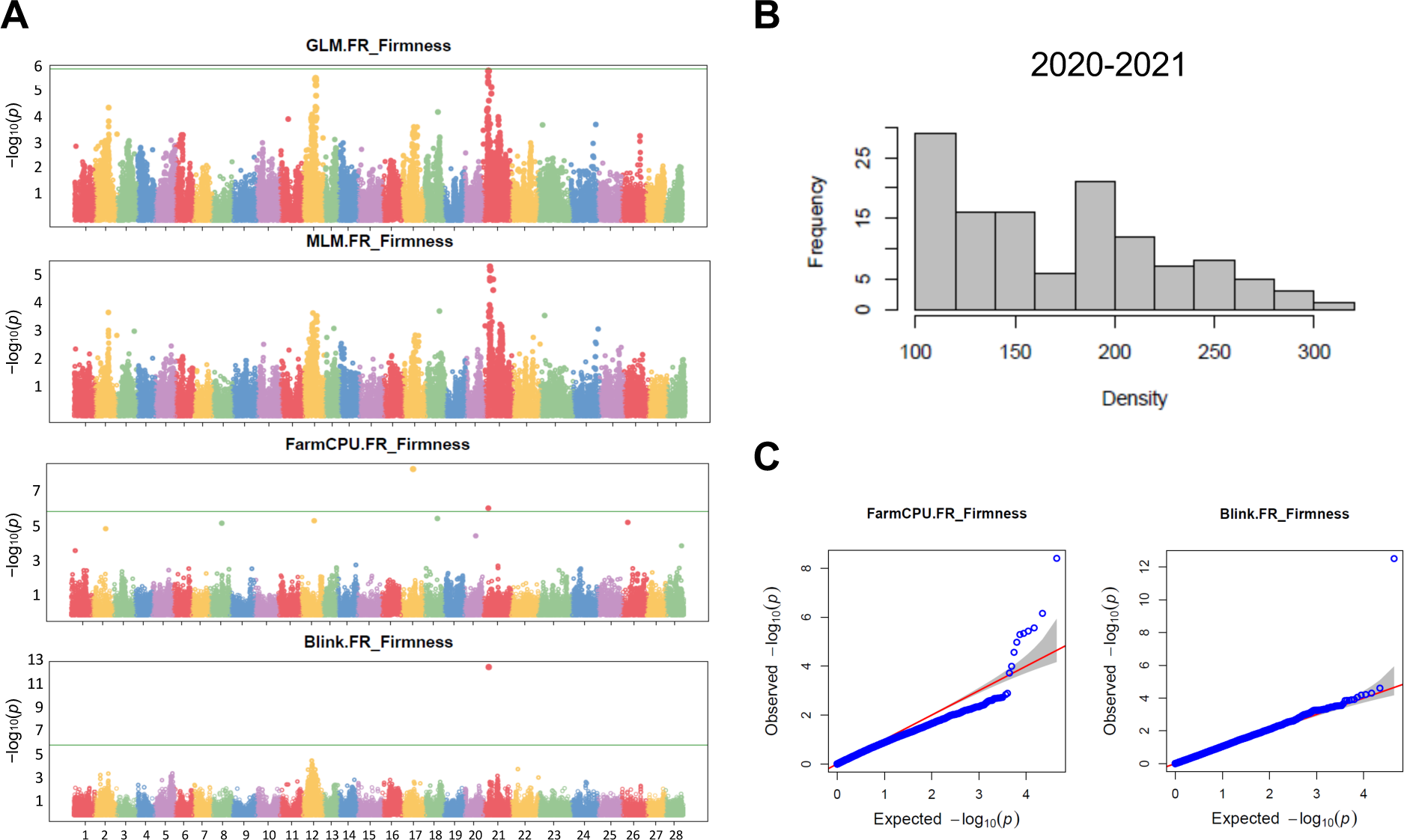
Significant GWAS associations for fruit firmness in 2020-2021. Manhattan plots of −log10P versus chromosomal position for the four models (A), phenotypic distribution of fruit firmness (B), and QQ plots of the model(s) with the highest number of associations (C). The plots of the different chromosomes are shown in different colors, which follow the order of ’Camarosa’ v.1.0 (Edger et al., 2019): chromosome 1 (1-1) to chromosome 28 (7-4). Solid and dashed green horizontal lines represent the significance thresholds following the Bonferroni correction method (-log10 (0.01/total SNPs) and (-log10(0.05/total SNPs), respectively).

### LD haploblock and candidate genes for fruit firmness QTL on chromosome 6A

As firmness is an important breeding target, and due to the large effect of the QTL for FR_Firmness in chromosome 6A, we decided to focus our search for the underlying gene on this region. Besides, a QTL for firmness in the same region has been recently detected in other GWAS using wild and domesticated cultivars (Hardigan et al., 2021a) and a multiparent population (Cockerton et al., 2021), suggesting that it is conserved in diverse genotypes and environments. The majority of significant SNPs detected on 6A in our study spanned a chromosomal region of 0.89 Mb in high linkage disequilibrium flanked by markers AX- 166518257 and AX-184933268 (Supplementary Table S4; Supplementary Fig. S12). This region contains 177 annotated genes in the ‘Camarosa’ reference genome, including four closely located genes with similarity to polygalacturonases (PGs; FxaC_21g15450, FxaC_21g15750, FxaC_21g15770 and FxaC_21g15780) and one transcript with similarity to endoglucanases (FxaC_21g15730; Supplementary Table S6). Fruit softening during ripening is caused by disassembly of the cell wall and the dissolution of the middle lamella, and a set of enzymes including polygalacturonases mediates this process (Moya-Leon et al., 2019). Therefore, these genes are good candidates for controlling the observed natural variation on fruit firmness in the GWAS population.

We next studied the expression of these five genes during ‘Camarosa’ fruit ripening using RNAseq data from a previous study (Sánchez-Sevilla et al., 2017) but here mapped to either the ‘Camarosa’ v1.0 a.2 (Edger et al., 2019) or the ‘Royal Royce’ FaRR1 (Hardigan et al., 2021b) genomes. Only one of them (FxaC_21g15770) showed significant expression in the analyzed tissues (Supplementary Table S7). Interestingly, this transcript encodes for the functionally characterized *FaPG1* gene (AF380299; Quesada et al., 2009). As in previous reports, in our RNAseq analysis *FaPG1* was induced during fruit ripening, with the highest expression in the turning and ripe stages of the receptacle tissue. While the gene annotation of *FaPG1* in ‘Camarosa’ (FxaC_21g15770) contains the four expected exons, in the FaRR1 reference genome its annotation (Fxa6Ag103973) is not correct. The Fxa6Ag103973 gene spans eight predicted exons from two incomplete tandemly arranged PG genes. However, the two complete open reading frames of these two genes, *FaPG1* and the next PG (FxaC_21g15750 in ‘Camarosa’) could be manually found in the ‘Royal Royce’ reference genome. An alignment of the four PG sequences from ‘Camarosa’ and ‘Royal Royce’ is shown in Supplementary Fig. S13. The gene *FaPG1* was identical in ‘Camarosa’ and ‘Royal Royce’. Similarly, high similarity was observed between the ‘Camarosa’ and ‘Royal Royce’ alleles for the other three *PGs*, which were not significantly expressed in the analyzed tissues.

As *FaPG1* was the only candidate gene being expressed, and to confirm its role on natural variation on fruit firmness, we quantified its expression levels by qRT-PCR in ripe fruit from two pools of accessions contrasting on fruit firmness (Fig. 7). Expression of *FaPG1* was 13- fold higher in the accessions with lower firmness, indicating that this is the most plausible cause of the observed variation in fruit firmness.

**Fig. 7.**
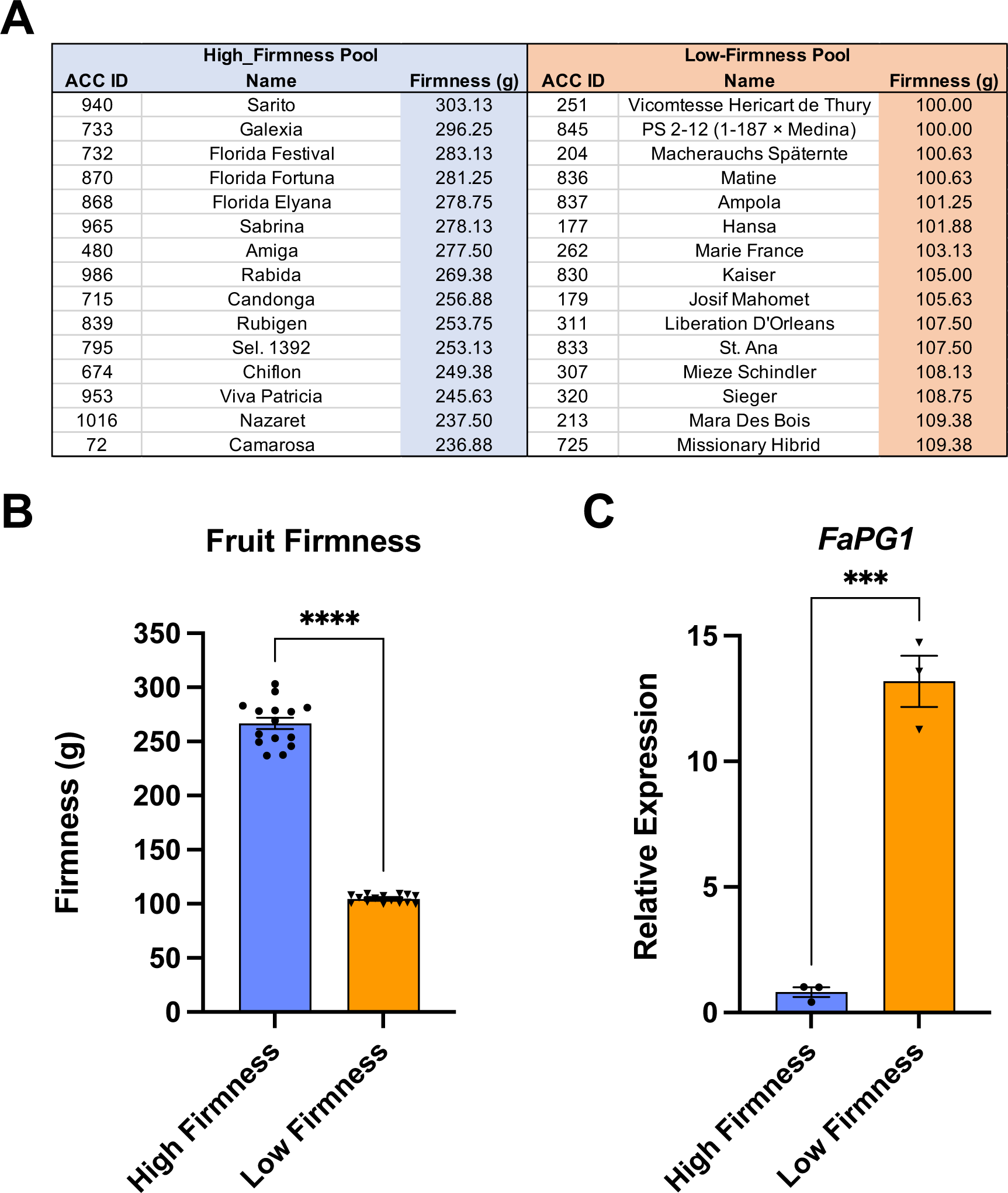
Fruit firmness and *FaPG1* expression in contrasting strawberry accessions. (A) Strawberry accessions in the high- and low-firmness pools. (B) Difference in fruit firmness between pools. (C) Relative expression of *FaPG1* by qRT-PCR in the two pools. Statistical significance by t test is shown (***p-value<0.005; ****p-value<0.001).

### Development and validation of a KASP assay for prediction of fruit firmness

Significant SNPs associated with the QTL on 6A were tested for KASP designing and resulted in a subgenome-specific assay for Axiom SNP AX-184242253. The predictive capacity of KASP-184242253 was tested on a collection of 138 strawberry accessions evaluated for fruit firmness during an additional fourth season (2022-2023; Supplementary Table S8). TT, CT and CC genotypic classes showed significant differences in fruit firmness (Fig. 8). Strawberry accessions with the homozygous reference allele (CC) produced fruits 47.56% firmer than those with the alternative TT allele. Heterozygous genotypes displayed an intermediate phenotype. A GWAS for fruit firmness using the 138 accessions and the 2022-2023 phenotypic data resulted once more in the detection of the same QTL on 6A (Supplementary Fig. S14). The phenotypic variance explained by this QTL was 44%. Together, these results indicate the great utility of this KASP assay for MAS.

**Fig. 8.**
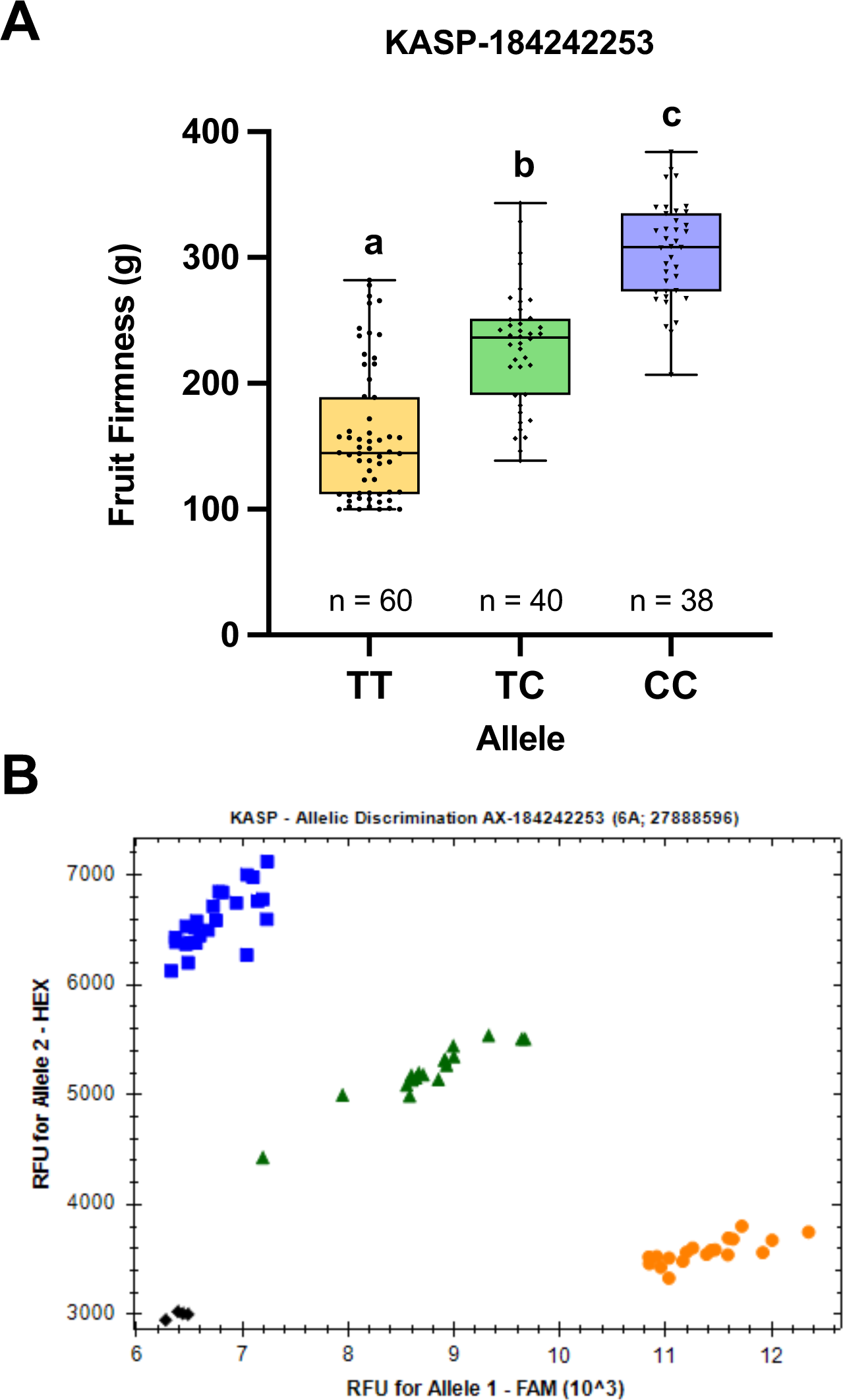
Development of a Kompetitive Allele Specific PCR (KASP) assay for fruit firmness prediction. (A) Allele effects of KASP-184242253 marker on fruit firmness in a collection of 138 accessions evaluated in season 2022-2023. (B) Example of genotype clusters for KASP- 184242253 marker. The reference C allele is labeled with HEX and the alternative T allele is labeled with FAM. Heterozygous samples are represented in green and non-template controls in black. Boxes span the 25th and 75th percentiles and the middle line represents the median. Whiskers (T-bars) are the minimum and maximum values. Letters show significant differences by one way ANOVA and Tukey test (*: p-value < 0.0001). n, number of accessions.

## Discussion

### Analysis of genetic diversity and population structure showed a diversification into six subpopulations, with considerable admixture and minor structuring

Population structure and PCA indicated a diversification based on the breeding history and the geographic adaptation, as in other reported analyses (Gil-Ariza et al., 2009; Zurn et al., 2022). Six subpopulations were identified, as in another recent study involving 1,300 octoploid strawberry accessions of both cultivated and wild species (Hardigan et al., 2021a). Older European varieties and the more recent Californian-Mediterranean type varieties (subpopulation 1 and 3 in Fig. 1C) were clearly separated, while three intermediate subpopulations contained much more genetic admixture (subpopulations 4, 5, and 2 in Fig. 1C). Both PCA and NJ tree separated the recent hybrids with *F. chiloensis* (Fig. 1A and C). Despite these results, the low FST value suggests that the different subpopulations still share considerable genetic diversity, and thus the collection is not very structured, which reduces the probability of false positives in GWAS. The value obtained for our population was similar to that reported in other studies using other octoploid populations (Hardigan et al., 2021a; Zurn et al., 2022).

### Relationship between traits and PCA highlight the main factors affecting phenotypic variation in the collection

In the three seasons, a large phenotypic variation was observed among individuals for all the evaluated traits, which was generally consistent among the three years (Table 1). According to this, positive correlations between seasons were in general medium-high. Correspondingly, phenotypic variation was in many traits also affected by different environmental factors between seasons. Correlations and PCA of phenotype data highlighted interesting relationships between traits (Fig. 2; Supplementary Table S3). First, the most vigorous genotypes produced the larger fruits, and the higher number of fruits (therefore, greater yields) and also higher number of runners. Similar conclusions were recently reported for another strawberry germplasm collection (Hummer et al., 2022). Our study also highlights a positive relationship between those traits and firmer fruits. In fact, fruit weight and firmness had a strong impact on the phenotypic variation observed among accessions and across the strawberry breeding history (Fig. 2B and C; Supplementary Fig. S1). In agreement, genomic signatures of domestication have been recently reported for these two breeding targets (Hardigan et al., 2021a; Fan and Whitaker, 2023).

Earlier runnering date correlated with greater production of runners. Also, more vigorous plants seemed to runner earlier. Negative correlations were also observed between runnering time and fruit number, indicating competition between vegetative and reproductive growth for available meristems (Gaston et al., 2013; Muñoz-Avila et al., 2022). Positive correlations between plant diameter and/or height with the number of runners have been previously reported (Antanaviciute et al., 2017; Hummer et al., 2022).

Fruit color traits were all significantly correlated and contributed greatly to the large diversity observed in the population (Fig. 2; Supplementary Table S3). As in other works (Zorrilla-Fontanesi et al., 2011; Lerceteau-Köhler et al., 2012; Ali & Serçe, 2022), we observed a strong positive correlation between fruit external lightness and yellow external hue. Both the external color and the internal color of the pulp (including the core) were positively correlated, which has been reported in other works, and is to be expected since all three depend on an accumulation of anthocyanins (Castillejo et al., 2020).

Strawberry accessions with higher SSC (FR_Brix) had fruits with higher vitamin C content (Fig. 2; Supplementary Table S3) in agreement with other reports (Zorrilla-Fontanesi et al., 2011; Ali & Serçe, 2022). Strikingly, these accessions belong mostly to the old European and American varieties (subpopulation 3), suggesting that breeding had a negative impact on these two important traits. In agreement with this conclusion, SSC and AsA were negatively correlated with fruit weight and fruit number, the main breeding targets throughout history.

Negative correlations between these traits have also been reported in other strawberry populations (Zorrilla-Fontanesi et al., 2011; Whitaker et al., 2012; Cockerton et al., 2021; Hummer et al., 2022). As previously reported, positive correlations were observed also between SSC and fruit acidity (Vallarino et al., 2019; Wada et al., 2020).

### Genotype-phenotype associations varied depending on the season and the GWAS model

Our GWAS for agronomic and fruit quality traits has detected 121 significant SNPs associated with 19 of the 26 evaluated traits, many of them represent novel and distinct QTL, while others overlap previously identified regions. Previously identified regions, as for fruit color traits on 1B caused by natural variation on *FaMYB10* (Castillejo et al., 2020), validate the experimental methods used, while novel associations, particularly those detected in more than one season and/or with larger phenotypic effects represent the starting point for the identification of the underlying genes. Also, significant markers with large effects here detected for many key breeding traits, such as runnering date and runner number, fruit weight or fruit firmness represent useful tools for MAS.

Association mapping is characterized by high resolution and power to detect monogenic traits, such as many resistance genes (Bush and Moore, 2012). However, for polygenic traits where many loci explain small amounts of phenotypic variation, the statistical signals are much weaker and more difficult to detect than for genetically simpler traits (Atwell et al., 2010). This might be the reason why for complex polygenic traits such as plant height, flower number, flowering time or AsA content, no significant associations were detected. Studying these complex traits would require a higher number of accessions (Balding, 2006; Bush and Moore, 2012), which may have limited the statistical power with our experimental population size. Twelve of the traits were evaluated with the use of scores, including FL_GCP, which was evaluated with a binomial scale. Although the use of scales might reduce the power of the association tests (Poland and Nelson, 2011), only in four of them, including FL_GCP, no significant associations were detected. However, petal greening was the only trait without significant correlations between any season, indicating that it might be mainly controlled by environmental factors.

Despite reasonable correlations between seasons and similar range of variation across years (Table 1), the majority of associations were detected in individual seasons, suggesting substantial environmental effects. GWAS results are not always consistent between years, as those reported by McClure et al. (2018) for fruit quality traits in apple, in which fruit firmness and color displayed high correlations between years, but different associations were detected for each year. In our study, most of the associations were detected in the third season (Table 3; Supplementary Table S4), which may have been influenced by the larger sample size. The use of four models allowed the evaluation of the best fit to our population, trying to avoid possible interactions that could produce false positives, or the lack of significant associations by false negatives. The BLINK model resulted in the highest number of significant SNPs, and in general produced QQ plots with the best connection between observed and expected p-values, indicating an effective control of population structure and family relatedness (Table 3; Supplementary Table S4).

### Differential expression of *FaPG1* controls natural variation in fruit firmness in strawberry

Improvement of fruit firmness has been a key breeding target since the origin of the species, about 300 years ago, due to its effects both on the organoleptic quality (texture) and postharvest and shelf life of the fruit. For this trait we identified seven QTL, although two of them, on chromosomes 3A and 6A, had larger phenotypic effects (Fig. 6; Supplementary Table S4). The QTL on chromosome 6A was detected in the third season, the one with the larger accession sample, using the four models and increased fruit firmness by about 27 g. Significant associations in exactly the same region have been detected in other populations (Hardigan et al., 2021a; Cockerton et al., 2021). We further detected this QTL using a larger sample size and the FaRR1 genome in an additional fourth season (Supplementary Fig. S14). The phenotypic variance explained (44%) and its detection in other populations and environments, indicate the usefulness of associated SNPs for MAS.

The 891 kb long haploblock enclosing the significant markers contains the *FaPG1* gene. Downregulation of *FaPG1* expression by antisense reduced strawberry fruit softening at ripe and postharvest stages (Garcia-Gago et al. 2009; Quesada et al. 2009), highlighting its key role on this trait. Further studies revealed a 42% decrease in pectin solubilization and reduced depolymerization of tightly bound pectins in *FaPG1*-antisense fruits (Posé et al. 2013). More recently, *FaPG1* knockout mutants generated by CRISPR/Cas9 editing resulted in fruits with increased firmness, reduced softening rate during postharvest, and less susceptibility to *Botritis cinerea* damage (López-Casado et al., 2023). We have shown a 13-fold change in *FaPG1* expression between strawberry accessions contrasting in fruit firmness (Fig. 7). We have also shown significant differences in fruit firmness across the six subpopulations of the germplasm collection (Supplementary Fig. S1), which agrees with the identification of the same QTL on 6A overlapping with early-phase domestication sweeps (Hardigan et al., 2021a). Finally, the KASP assay here developed was able to predict a 47.6% increase in fruit firmness, representing an extremely useful tool for MAS. Together, our results indicate that *FaPG1* is the main contributor to natural variation in strawberry fruit firmness.

### Supplementary data

Fig. S1. Phenotypic distribution of fruit firmness in the six different subpopulations of the GWAS.

Fig. S2. Significant GWAS associations for flower diameter. Fig. S3. Significant GWAS associations for plant diameter.

Fig. S4. Significant GWAS associations for value *b*\* of the fruit color.

Fig. S5. Significant GWAS associations for leaf color.

Fig. S6. Significant GWAS associations for fruit weight.

Fig. S7. Significant GWAS associations for value L* of the fruit color.

Fig. S8. Significant GWAS associations for red color on petals.

Fig. S9. Significant GWAS associations for fruit number.

Fig. S10. Significant GWAS associations for value a* of the fruit color.

Fig. S11. Significant GWAS associations for soluble solids content.

Fig. S12. Linkage disequilibrium (LD) heatmap of the haploblock of 6A QTL controlling fruit firmness.

Fig. S13. Alignment of polygalacturonases in the QTL for fruit firmness on 6A.

Fig. S14. GWAS for fruit firmness in the 2022-2023 season.

Table S1. Strawberry accessions used in this study.

Table S2. Genetic similarity between accessions.

Table S3. Correlations between traits in the 2020-2021 season.

Table S4. Significant SNPs detected in the GWAS.

Table S5. Gene annotation of the closest transcript to significant SNPs detected by GWAS.

Table S6. Predicted genes in the QTL for fruit firmness on chromosome 6A.

Table S7. Expression of candidate genes in ‘Camarosa’ tissues.

Table S8. Phenotype and genotype for fruit firmness in the season 2022-2023.

## Acknowledgements

1. P. Muñoz acknowledged a PhD FPI-INIA contract co-financed by FSE. We are grateful to Francisco J. Durán for his excellent care of strawberry plants.

## Author contributions

JFSS, and IA: conceptualization; PM, FJRG, and JFSS: data curation; PM, FJRG, NO, and MRV: investigation; PM, FJRG, CC, JFSS, and IA: formal analysis; CC: methodology; SV, and JFSS: software; PM: writing - original draft; IA: writing - review & editing; PM, FJRG, and IA: visualization; JFSS, CC, and IA: supervision; JFSS, and IA: funding acquisition.

## Data availability

All the data used in this study is provided in the Supplementary data.

## Abbreviations

ACC: Accession
AsA: Ascorbic Acid
CR: Call Rate
GWAS: Genome Wide Association Study
FST: Fixation index
IBS: Identity By State
KASP: Kompetitive Allele Specific PCR
PCA: Principal Component Analysis
PG: Polygalacturonase
QTL: Quantitative Trait Loci

