## Supplemental Figures for "Genome-wide association studies in a diverse strawberry collection unveil loci controlling agronomic and fruit quality traits"

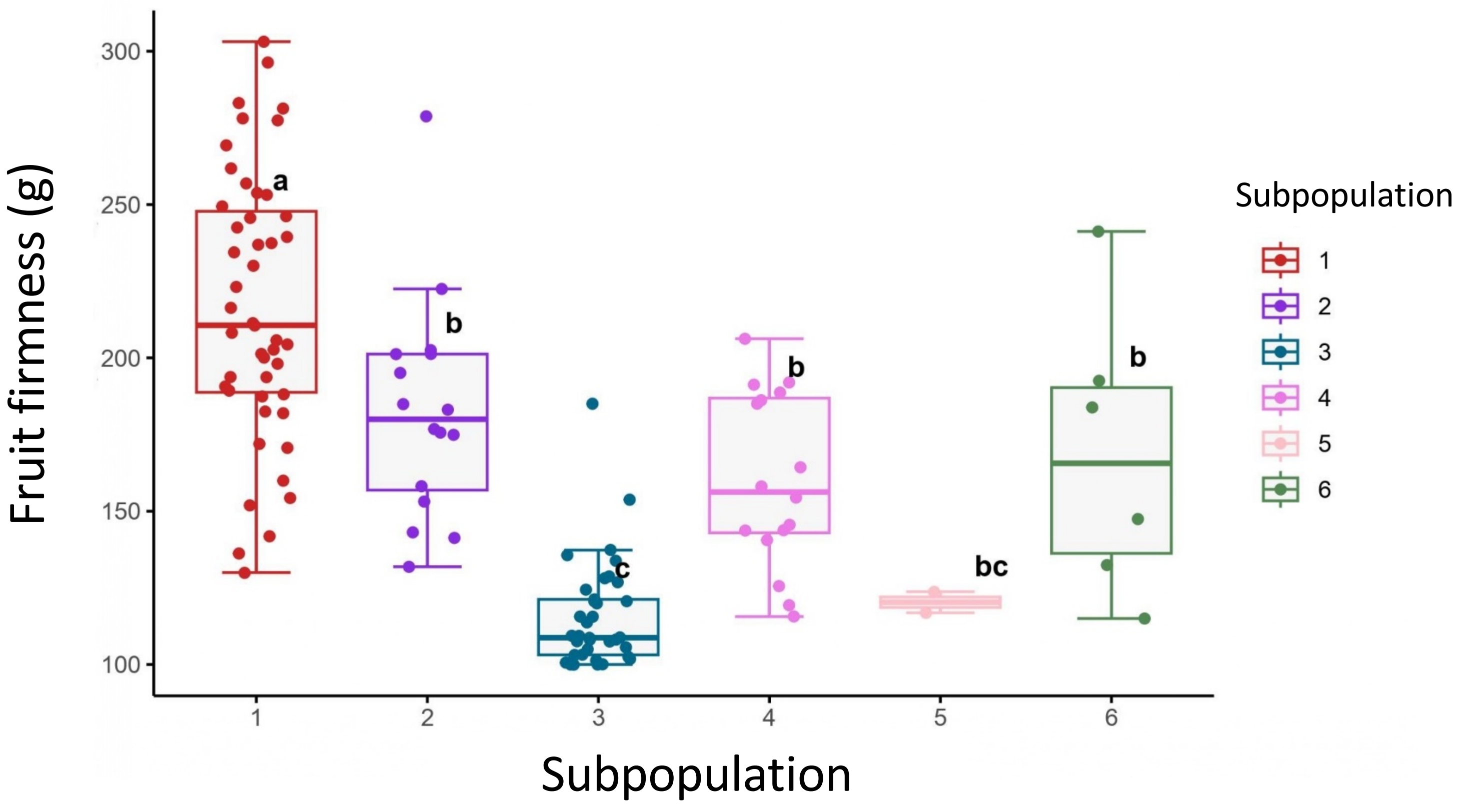


**Supplementary Fig. S1.** Phenotypic distribution of fruit firmness (g) in the six different subpopulations of the GWAS. The boxes includes the 25th and 75th percentiles and the solid bar within each box represent the median. T-bars represent confidence interval 95% and dots represent individual values. Each letter (*a, b* and *c*) indicates significant differences of $p < 0.05$ using ANOVA and Tukey-Kramer post hoc test for multiple comparisons.

### Flower diameter (2019-2020)

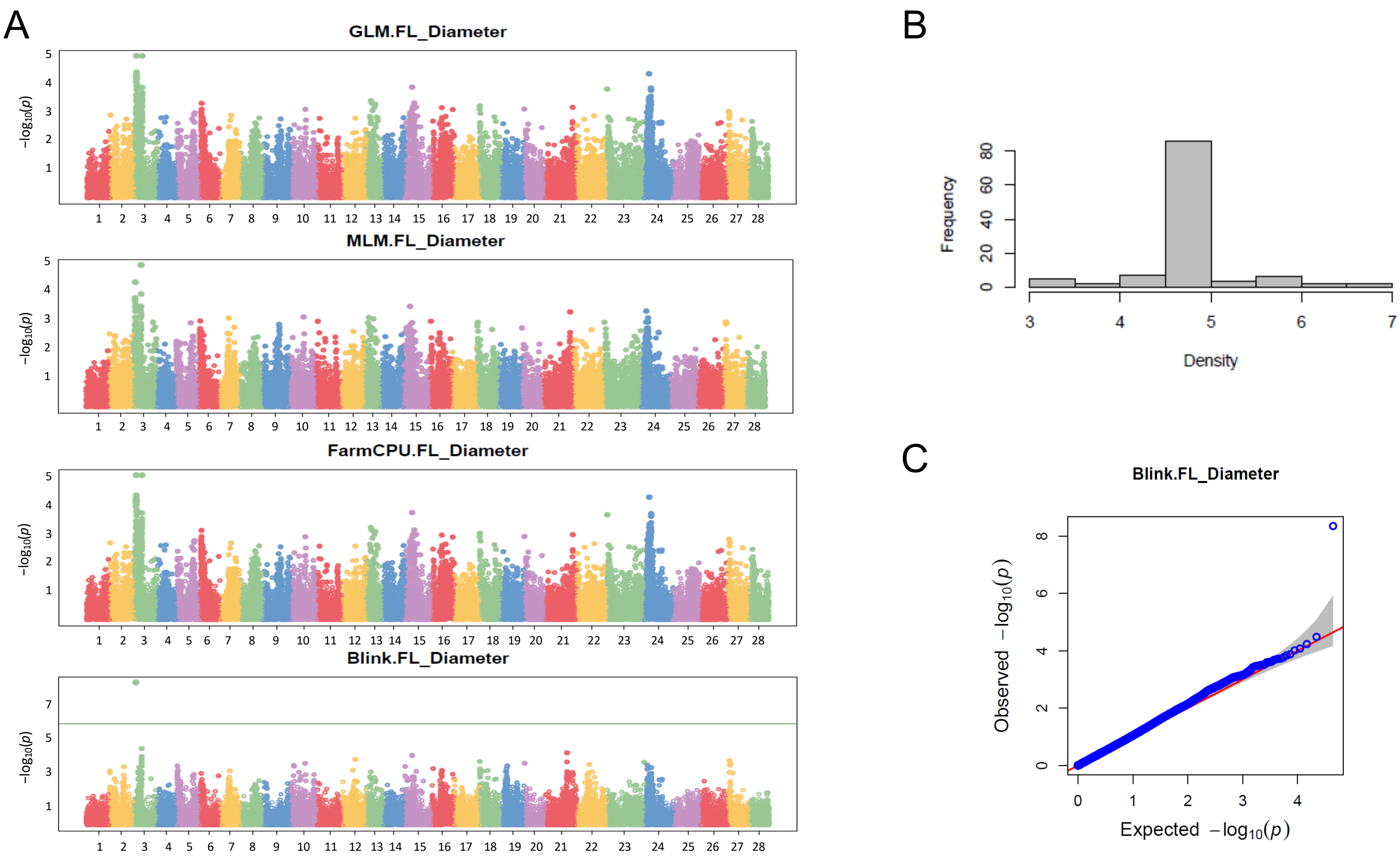


**Supplementary Fig. S2.** Significant GWAS associations for flower diameter. Manhattan plots of −log10 P versus chromosomal position for the four models (A) , phenotypic distribution of the trait (B), and QQ plots of the model with significant associations (C). Different chromosomes are shown in different colors, which follow the order of 'Camarosa' v.1.0 (Edger et al., 2019): chromosome 1 (1-1) to chromosome 28 (7-4). The green horizontal line represents the significance threshold using FDR.

#### Plant diameter (2020-2021)

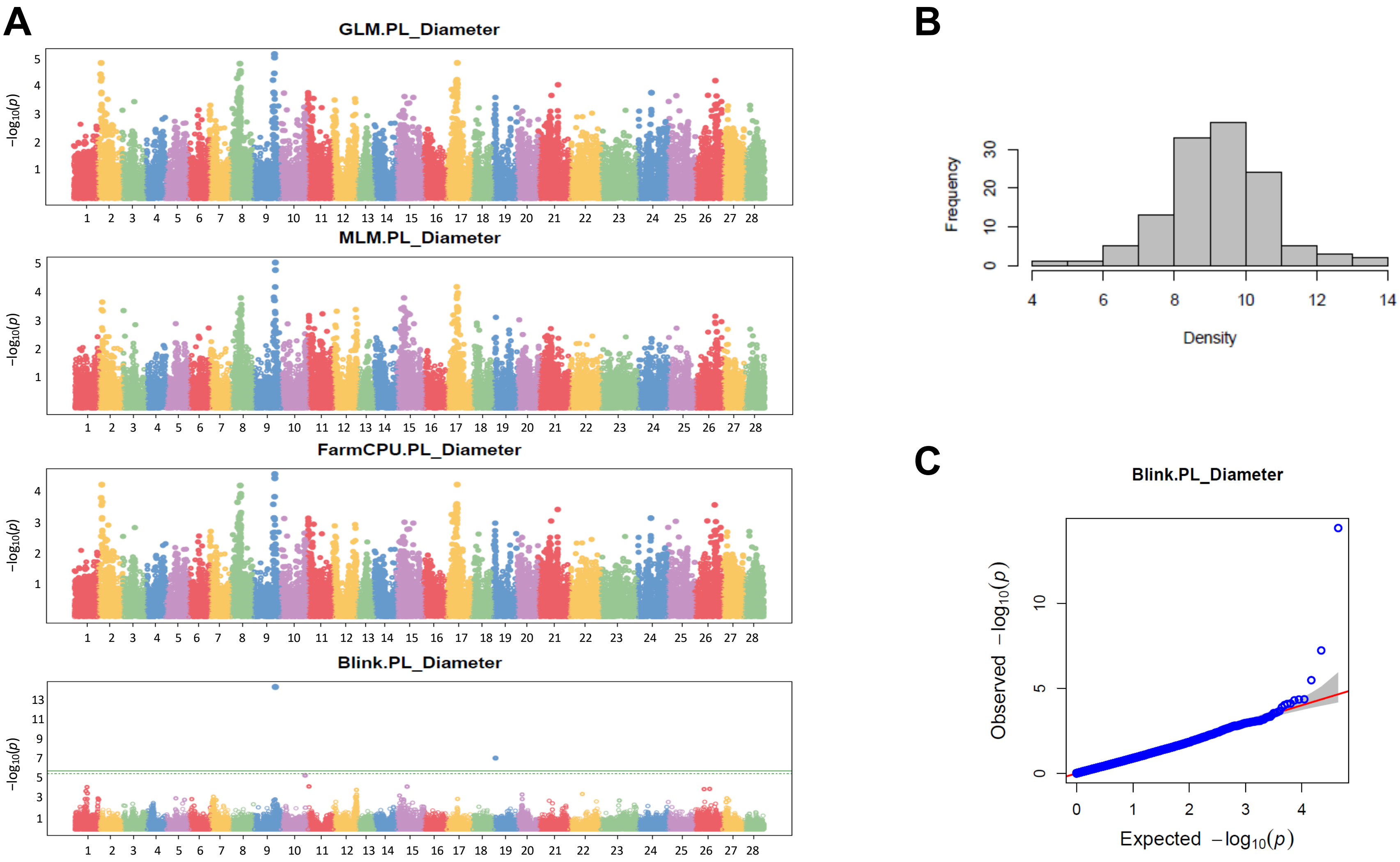


**Supplementary Fig. S3.** Significant GWAS associations for plant diameter. Manhattan plots of −log10 P versus chromosomal position for the four models (A), phenotypic distribution of the trait (B), and QQ plots of the model with significant associations (C). Different chromosomes are shown in different colors, which follow the order of 'Camarosa' v.1.0 (Edger et al., 2019): chromosome 1 (1-1) to chromosome 28 (7-4). The green horizontal line represents the significance threshold using FDR.

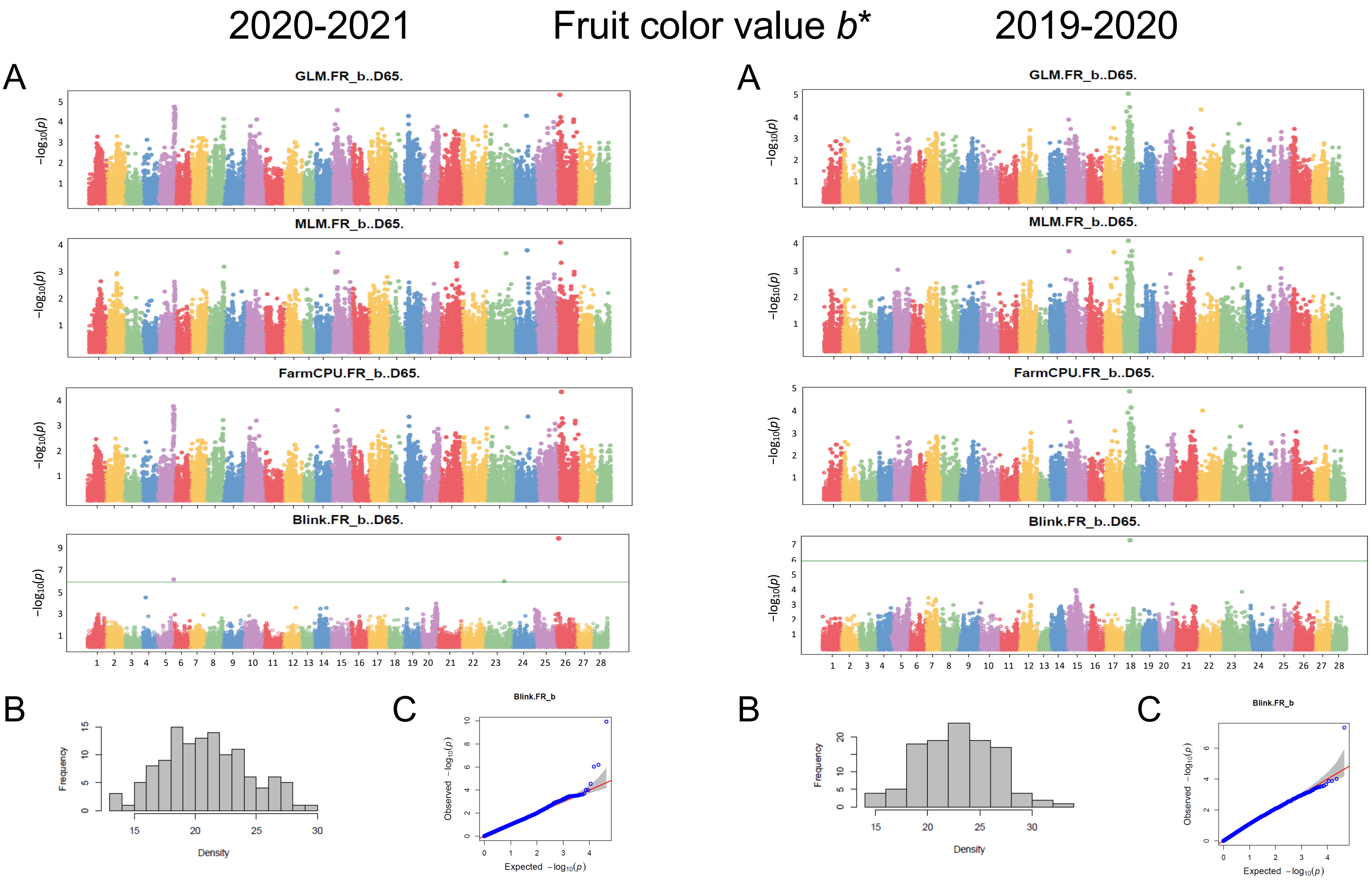


**Supplementary Fig. S4.** Significant GWAS associations for value *b** of the external fruit color. Manhattan plots of −log10 P versus chromosomal position for the four models (A), phenotypic distribution of the trait (B), and QQ plots of the model with significant associations (C). Different chromosomes are shown in different colors, which follow the order of 'Camarosa' v.1.0 (Edger et al., 2019): chromosome 1 (1-1) to chromosome 28 (7-4). The green horizontal line represents the significance threshold using FDR.

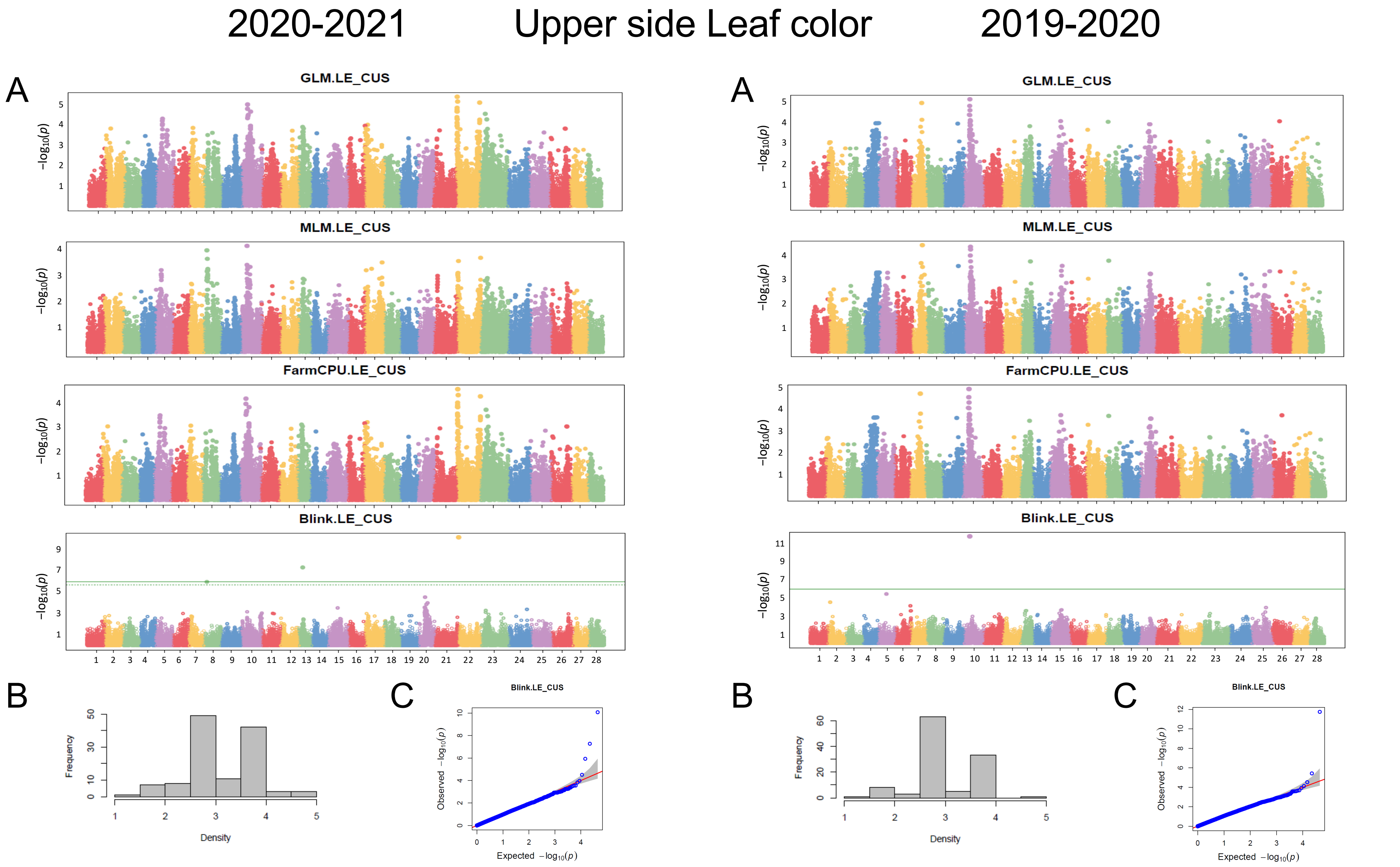


**Supplementary Fig. S5.** Significant GWAS associations for leaf color upper side. Manhattan plots of −log10 P versus chromosomal position for the four models (A), phenotypic distribution of the trait (B), and QQ plots of the model with significant associations (C). Different chromosomes are shown in different colors, which follow the order of 'Camarosa' v.1.0 (Edger et al., 2019): chromosome 1 (1-1) to chromosome 28 (7-4). The green horizontal line represents the significance threshold using FDR.

### Fruit weight (2020-2021)

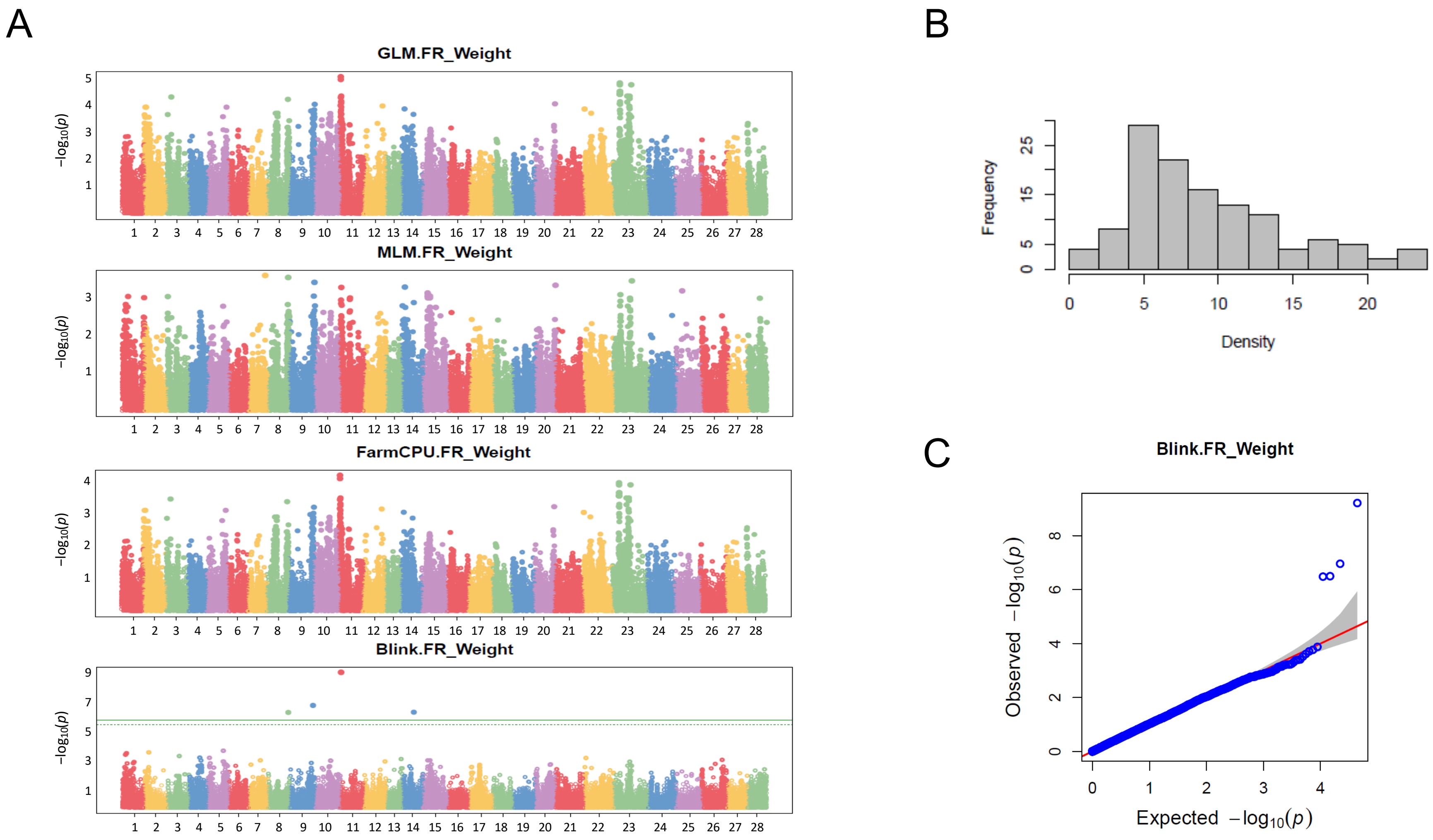


**Supplementary Fig. S6.** Significant GWAS associations for fruit weight. Manhattan plots of −log10 P versus chromosomal position for the four models (A), phenotypic distribution of the trait (B), and QQ plots of the model with significant associations (C). Different chromosomes are shown in different colors, which follow the order of 'Camarosa' v.1.0 (Edger et al., 2019): chromosome 1 (1-1) to chromosome 28 (7-4). The green horizontal line represents the significance threshold using FDR.

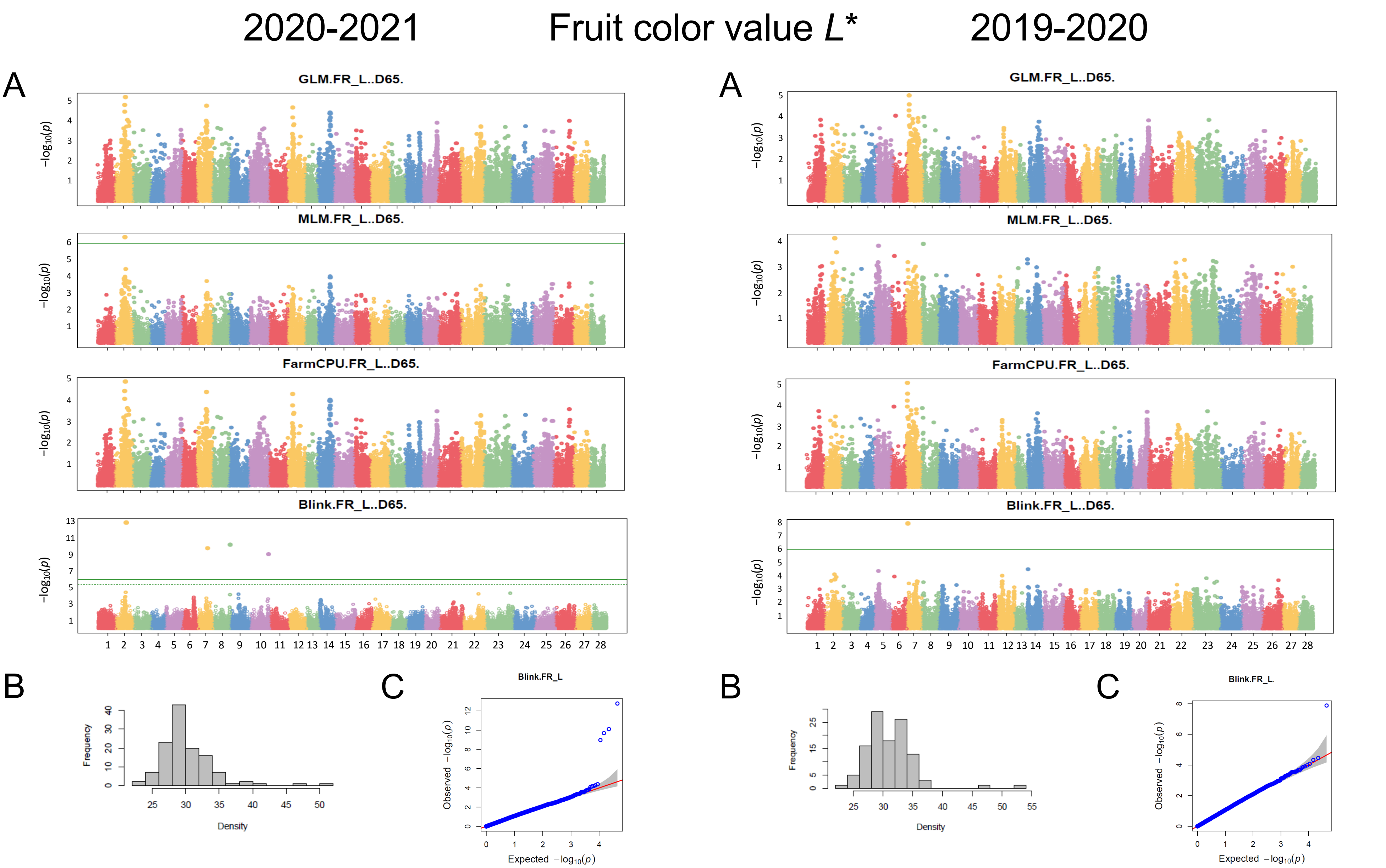


**Supplementary Fig. S7.** Significant GWAS associations for value *L** of the external fruit color. Manhattan plots of −log10 P versus chromosomal position for the four models (A), phenotypic distribution of the trait (B), and QQ plots of the model with significant associations (C). Different chromosomes are shown in different colors, which follow the order of 'Camarosa' v.1.0 (Edger et al., 2019): chromosome 1 (1-1) to chromosome 28 (7-4). The green horizontal line represents the significance threshold using FDR.

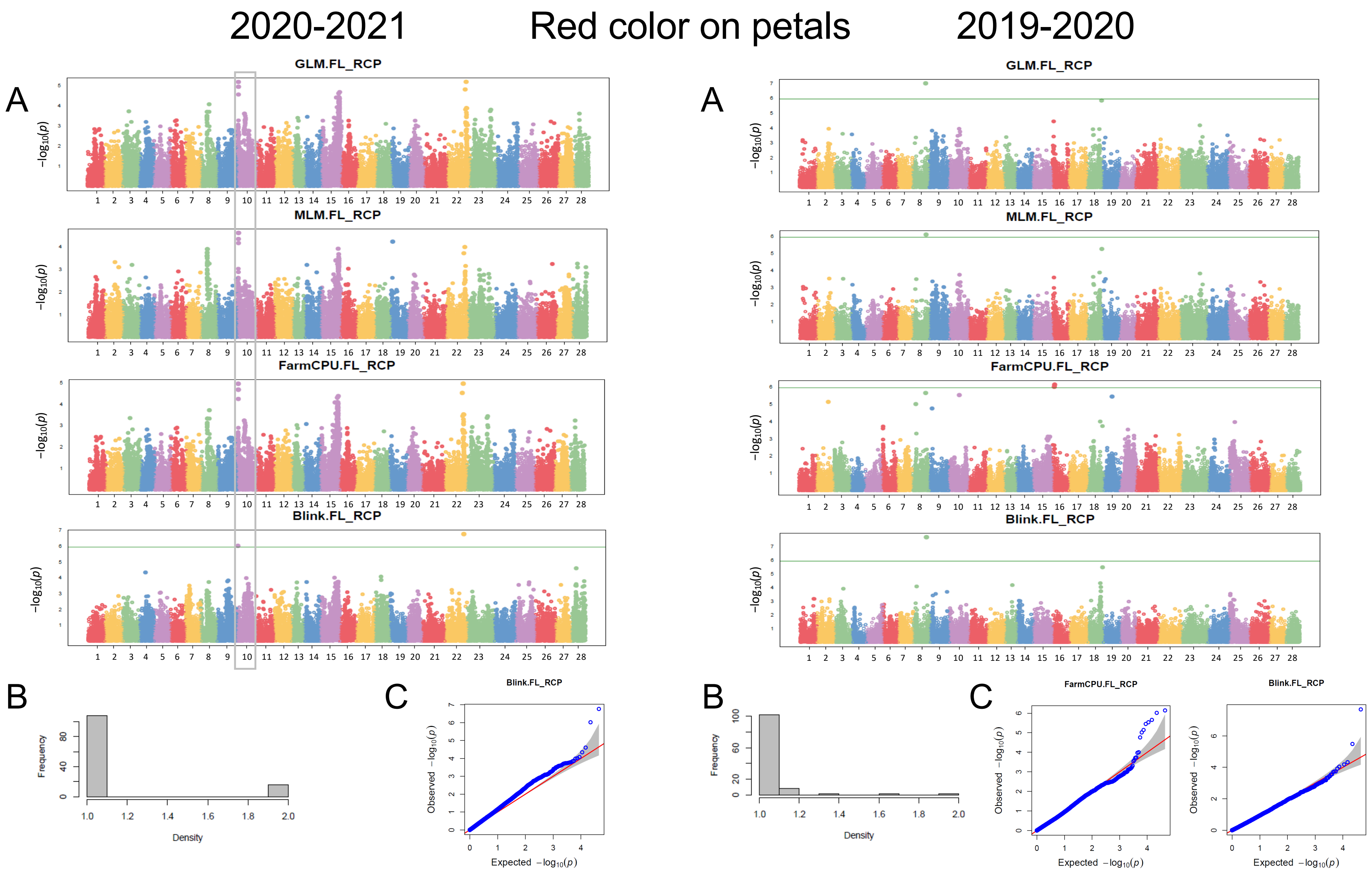


**Supplementary Fig. S8.** Significant GWAS associations for red color on petals. Manhattan plots of −log10 P versus chromosomal position for the four models (A) , phenotypic distribution of the trait (B), and QQ plots of the model with significant associations (C). Different chromosomes are shown in different colors, which follow the order of 'Camarosa' v.1.0 (Edger et al., 2019): chromosome 1 (1-1) to chromosome 28 (7-4). The green horizontal line represents the significance threshold using FDR. QTL detected in more than one season is highlighted.

### Red color on petals (2018-2019)

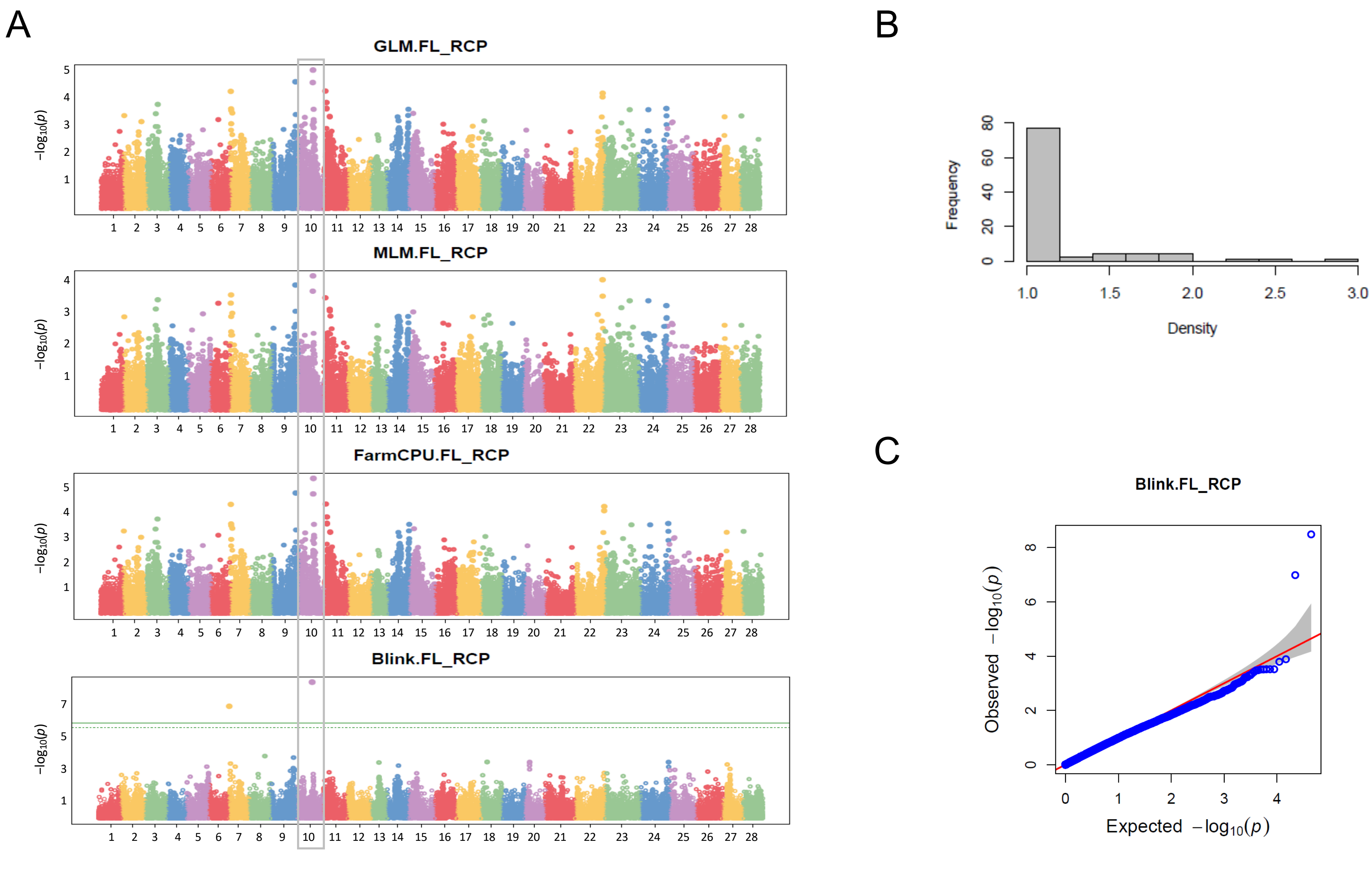


**Supplementary Fig. S8, continued.** Significant GWAS associations for Red Color on Petals. Manhattan plots of −log10 P versus chromosomal position for the four models (A), phenotypic distribution of the trait (B), and QQ plots of the model with significant associations (C). Different chromosomes are shown in different colors, which follow the order of 'Camarosa' v.1.0 (Edger et al., 2019): chromosome 1 (1-1) to chromosome 28 (7-4). The green horizontal line represents the significance threshold using FDR. QTL detected in more than one season is highlighted.

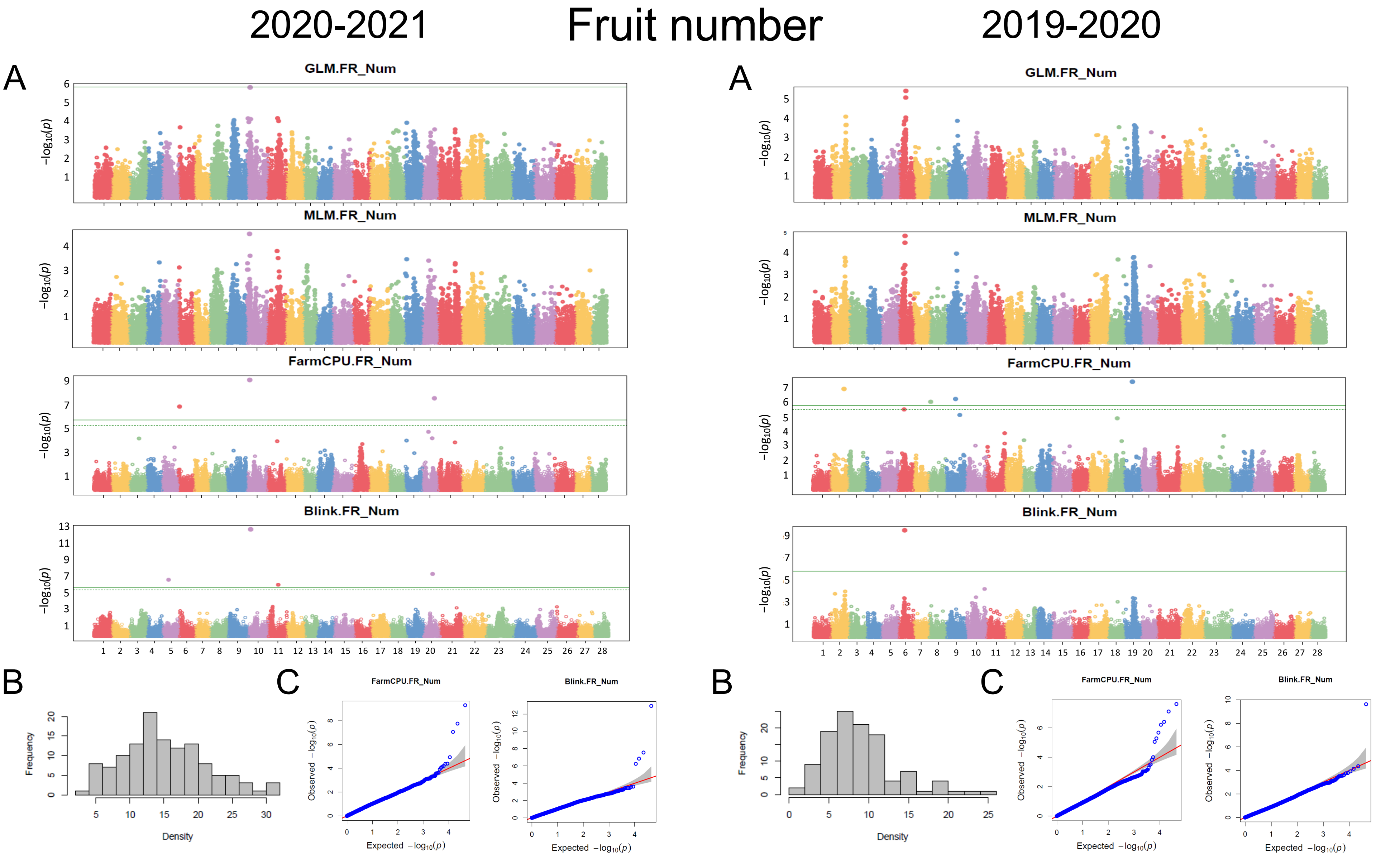


**Supplementary Fig. S9.** Significant GWAS associations for fruit number. Manhattan plots of −log10 P versus chromosomal position for the four models (A), phenotypic distribution of the trait (B), and QQ plots of the model with significant associations (C). Different chromosomes are shown in different colors, which follow the order of 'Camarosa' v.1.0 (Edger et al., 2019): chromosome 1 (1-1) to chromosome 28 (7-4). The green horizontal line represents the significance threshold using FDR.

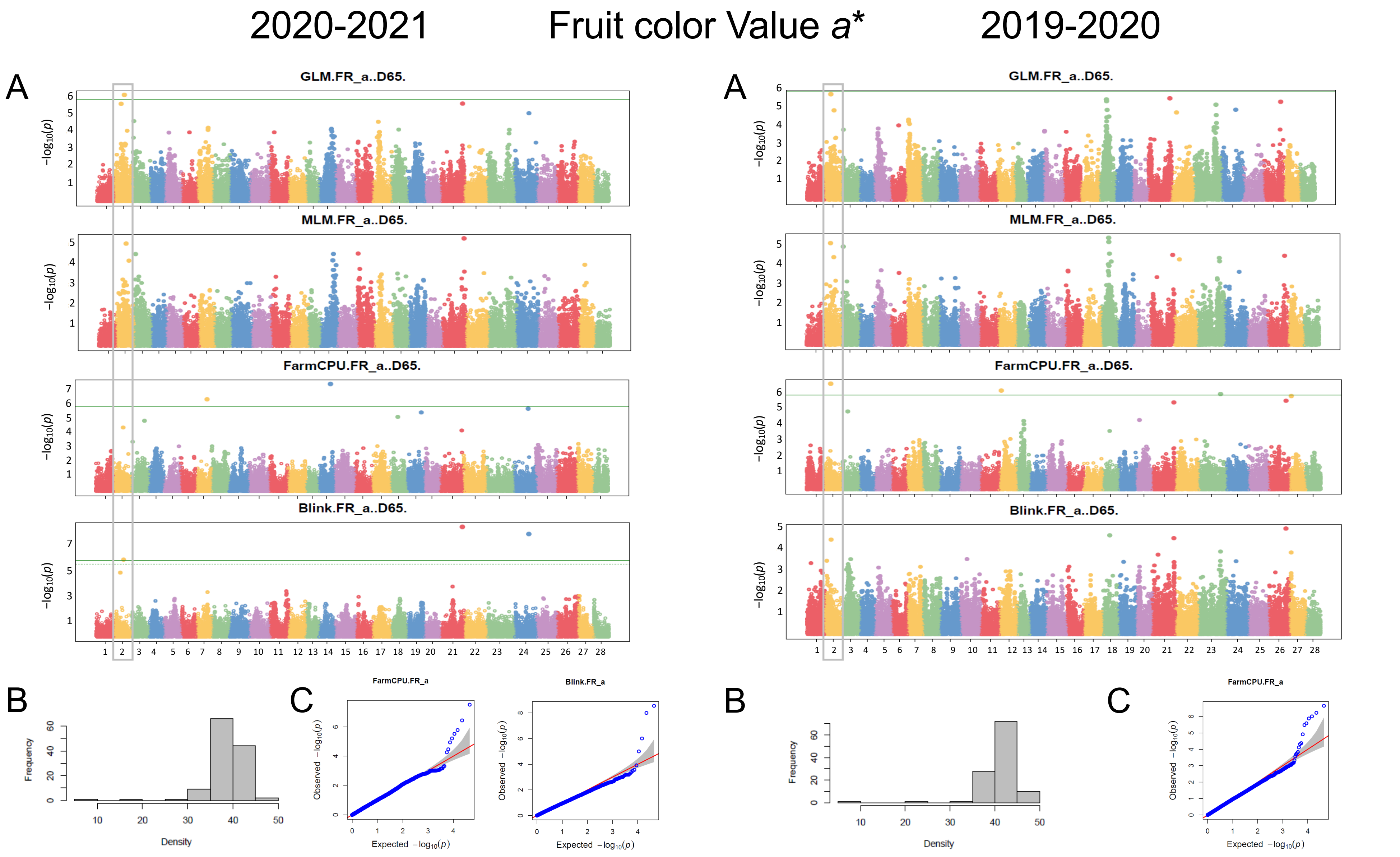


**Supplementary Fig. S10.** Significant GWAS associations for value *a** of the external fruit color. Manhattan plots of −log10 P versus chromosomal position for the four models (A), phenotypic distribution of the trait (B), and QQ plots of the model with significant associations (C). Different chromosomes are shown in different colors, which follow the order of 'Camarosa' v.1.0 (Edger et al., 2019): chromosome 1 (1-1) to chromosome 28 (7-4). The green horizontal line represents the significance threshold using FDR. QTL detected in more than one season is highlighted.

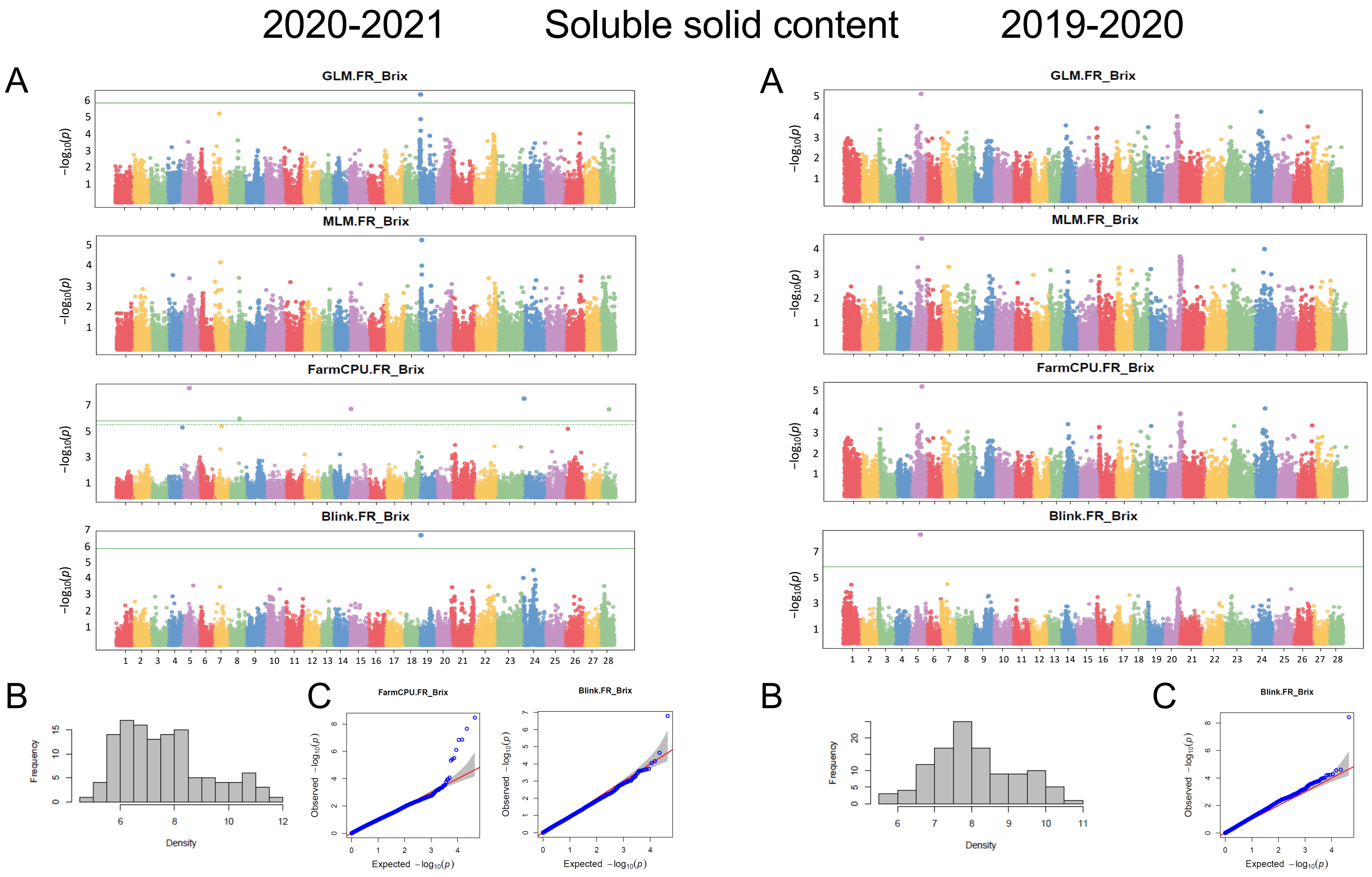


**Supplementary Fig. S11.** Significant GWAS associations for soluble solids content. Manhattan plots of −log10 P versus chromosomal position for the four models (A), phenotypic distribution of the trait (B), and QQ plots of the models with significant associations (C). Different chromosomes are shown in different colors, which follow the order of 'Camarosa' v.1.0 (Edger et al., 2019): chromosome 1 (1-1) to chromosome 28 (7-4). The green horizontal line represents the significance threshold using FDR.

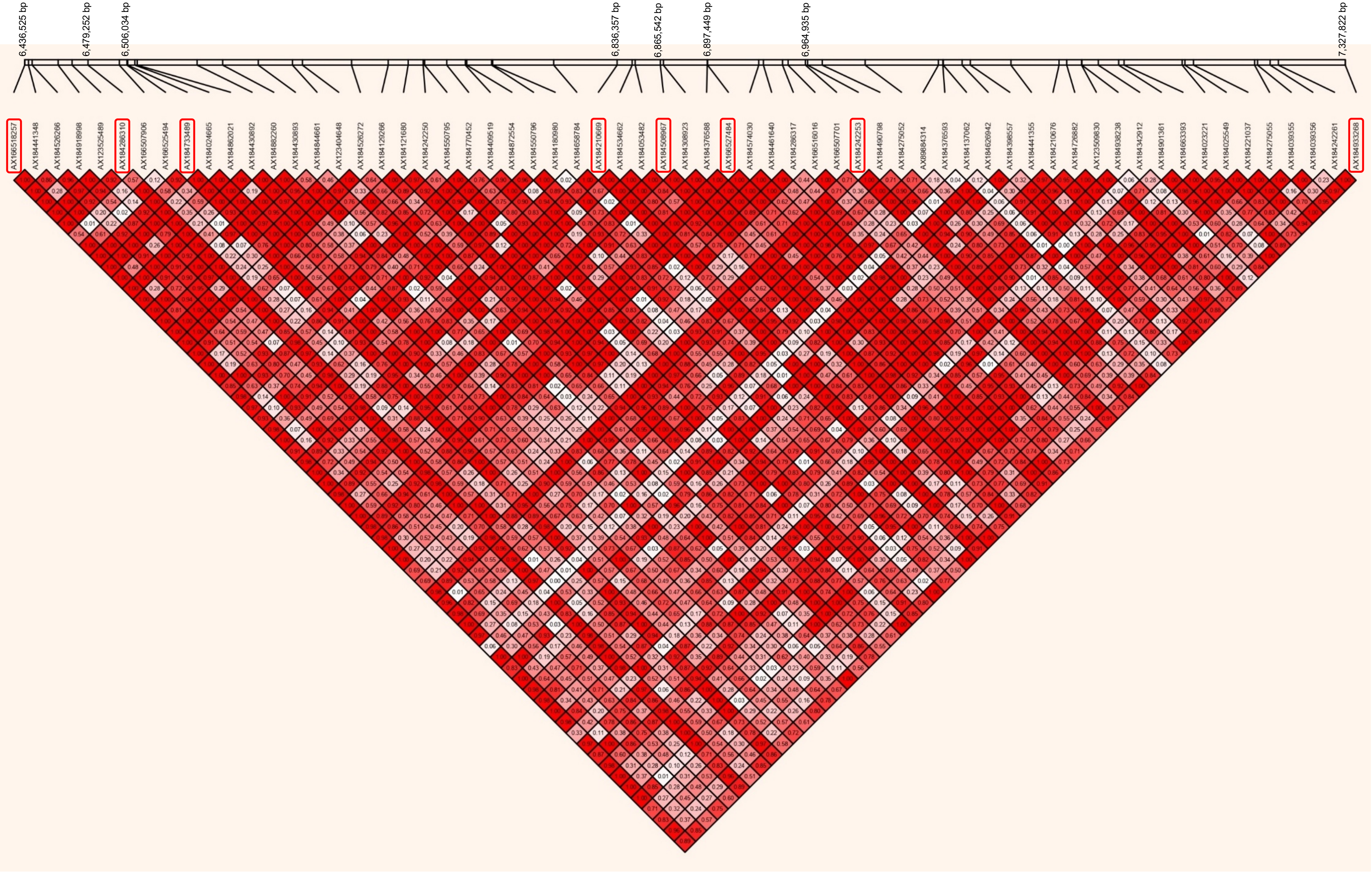


**Supplementary Fig. S12.** Linkage disequilibrium (LD) heatmap of the haploblock of 6A QTL controlling fruit firmness. The physical position of significant SNPs is given in the bar above the plot. The pairwise LD (D’) values are given in each box and represented by a scale from white (low) to dark-red (high). Significant SNPs detected by GWAS are highlighted. This region is 891 kb long and encloses 177 genes.

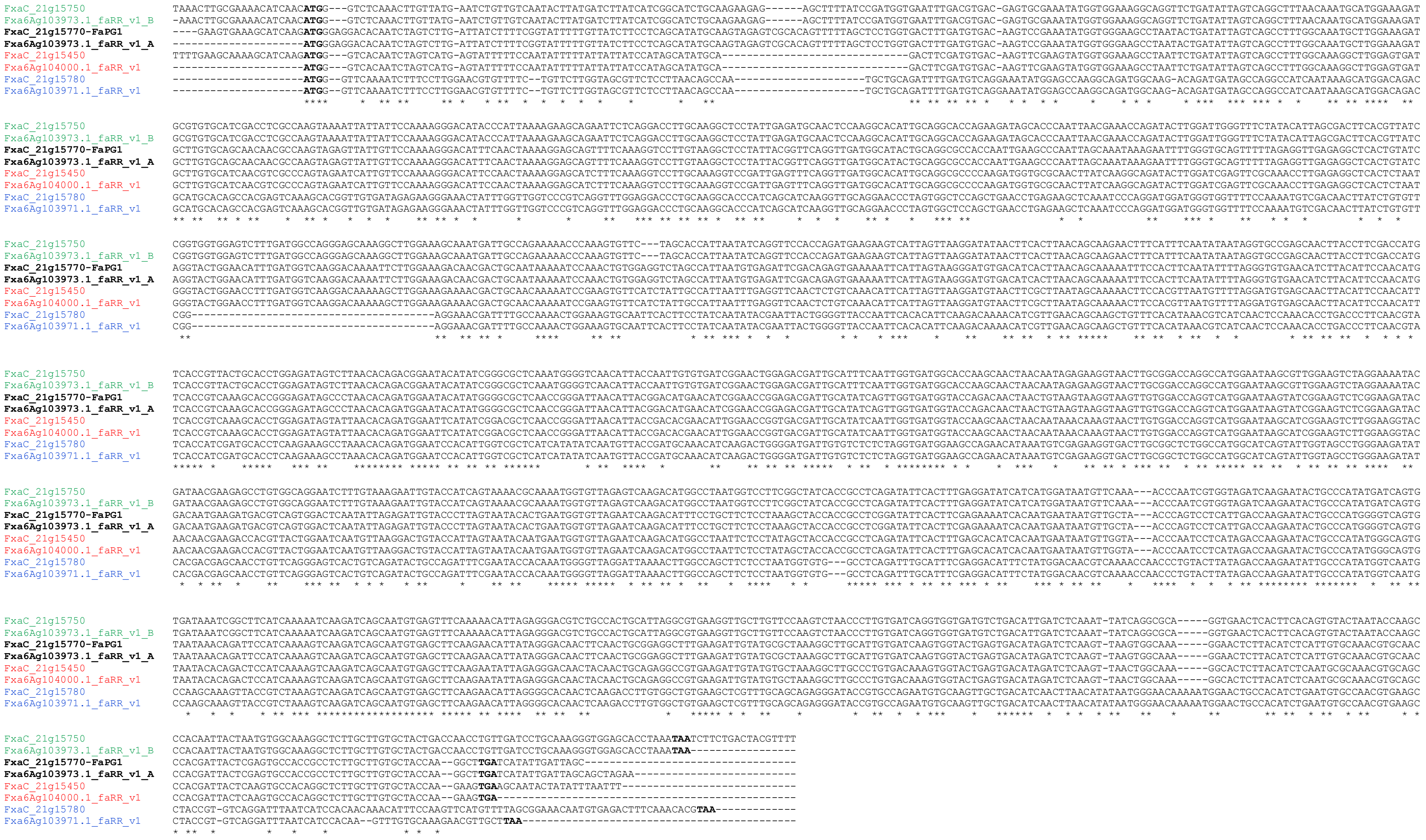


**Supplementary Fig. 13.** Clustal Omega alignment of the four *polygalacturonase* genes from 'Camarosa' and 'Royal Royce' in the QTL region for fruit firmness on 6A. Names of orthologous gene pairs are colored differentially. Fxa6Ag103973.1_faRR_v1_A and B have been manually extracted from FaRR1 genome as the gene annotation partially includes the two different *Polygalacturonase* genes.

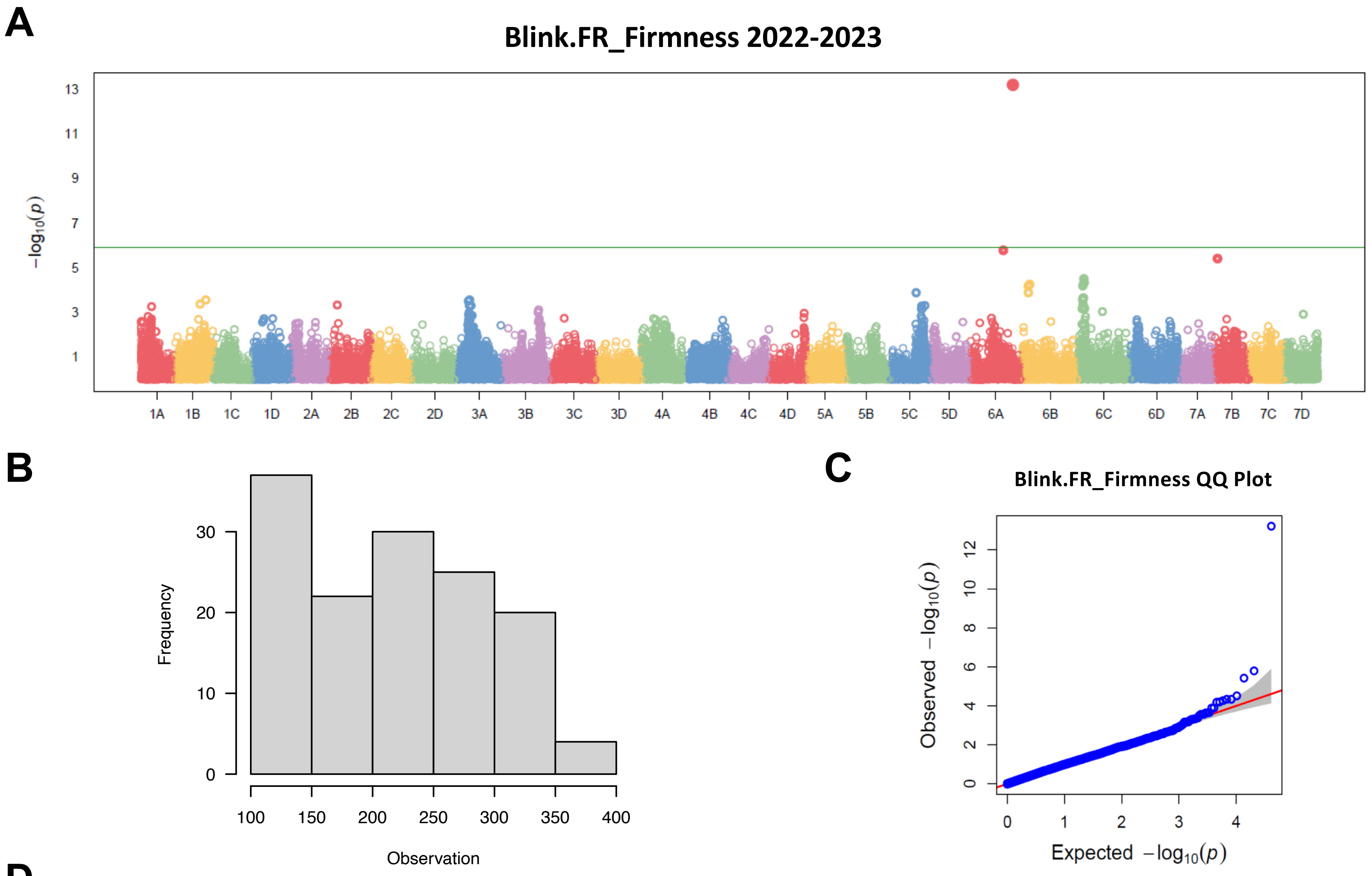


**D**

| SNP | Chr | Position | P.value | MAF | H&B.P.Value | Effect | PVE (%) |
|---|---|---|---|---|---|---|---|
| AX-184210669 | 6A | 28,017,174 | 6.19E-14 | 0.42 | 2.53E-09 | -43.39 | 43.97 |

**Supplementary Fig. S14.** Significant GWAS associations for fruit firmness in the 2022-2023 season. Manhattan plots of −log10 P versus chromosomal position for the BLINK model (A), phenotypic distribution of the trait (B), QQ plot (C), and details of the significant SNP on 6A (D). Different chromosomes are shown in different colors, which correspond to the 'Royal Royce' FaRR1 reference genome (Hardigan et al., 2021). The green horizontal line represents the significance threshold using FDR.
